# Subcellular localization of PKA catalytic subunits provides a basis for their distinct functions in the retina

**DOI:** 10.1101/2021.05.17.444584

**Authors:** Jinae N. Roa, Yuliang Ma, Zbigniew Mikulski, Qianlan Xu, Ronit Ilouz, Susan S. Taylor, Dorota Skowronska-Krawczyk

**Author notes:** **Corresponding author:** Susan S. Taylor, Dorota Skowronska-Krawczyk.

## Abstract

PKA signaling is essential for numerous processes but the subcellular localization of specific PKA isoforms has yet to be explored comprehensively in tissues. Expression of the Cβ protein, in particular, has not been mapped previously at the tissue level. In this study we used retina as a window into PKA signaling in the brain and characterized localization of PKA Cα, Cβ, RIIα, and RIIβ subunits. Each subunit presented a distinct localization pattern. Cα and Cβ were localized in all tissue layers, while RIIα and RIIβ were enriched in the photoreceptor cells in contrast to the cell body and retinal portion of retinal ganglion cells. Only Cα was observed in photoreceptor outer segments and the cilia transition zone, while Cβ was localized primarily to mitochondria and was especially prominent in the ellipsoid of the cone cells. In contrast to Cα, Cβ also never colocalized with RIIα or RIIβ. Using BaseScope technology to track expression of the Cβ isoforms we find that Cβ4 and Cβ4ab are prominently expressed and, therefore, likely code for mitochondrial-Cβ proteins. Our data indicates that PKA subunits are functionally nonredundant in the retina and suggesting that Cβ might be important for mitochondrial-associated neurodegenerative diseases previously linked to PKA dysfunction.

## Introduction

Vision is the most appreciated of the five senses, and age- and disease-related loss of visual perception has devastating impacts on quality of life. Several studies with mouse retina degeneration models implicate the second messenger cyclic adenosine 3’,5’ monophosphate (cAMP) and cAMP-dependent protein kinase A (PKA). Early works indicate that the maintenance of tightly regulated intracellular levels of cAMP plays a critical role in the modulation of photoreceptor light adaptation (Cohen and Blazynski, 1990; Cohen et al., 1992; Nir et al., 2002) and rod outer segment shedding and renewal (Stenkamp et al., 1994); and recent studies using PKAchu mice suggest a role for PKA in the dark adaptation in rods (Sato et al., 2020). PKA signaling at mitochondria is also important for fission/fusion and mitophagy (Dagda and Banerjee, 2015), both of which are essential for maintaining high rates of oxidative phosphorylation in the photoreactive zones of the retina. No study to date, however, has focused specifically on the role of subunit specific PKA signaling in the retina and, in particular, on the role of the Cβ isoforms.

Eukaryotic cells express multiple forms of PKA regulatory (R) and catalytic (C) subunits, and this subunit diversity accounts for a large part of its functional specificity. In general, PKA holoenzyme consists of an R subunit dimer bound to two C subunits (R_2_C_2_) (**Figure 1A**). The biochemical and functional features of PKA holoenzymes are largely determined by the structure and the biochemical properties of the four functionally non-redundant regulatory subunits, RIα, RIβ, RIIα and RIIβ (Mcknight et al., 1988; Ilouz et al., 2012; Taylor et al., 2012; Zhang et al., 2012). Spatially restricted localization of these holoenzymes, mediated primarily by scaffold proteins, referred to as A Kinase Anchoring proteins (AKAPs), provides an important layer of specificity in PKA signaling as was demonstrated by the unique expression of RIβ and RIIβ in brain (Ilouz et al., 2017). Although current dogma emphasizes the importance of subcellular localization of AKAP-bound R subunits in the functional diversity of PKA signaling (Pawson and Scott, 2010), little attention has been paid to the isoform diversity of the C subunits. Interestingly, while Cα is ubiquitously expressed in all mammalian cells, RNA *in situ* hybridization shows 50% of PKA signaling in brain is likely due to Cβ (Ørstavik et al., 2001), but without spatial information on endogenous protein expression, it is difficult to hypothesize how these two C subunits might influence regulation of specific neuronal processes.

**Figure 1.**
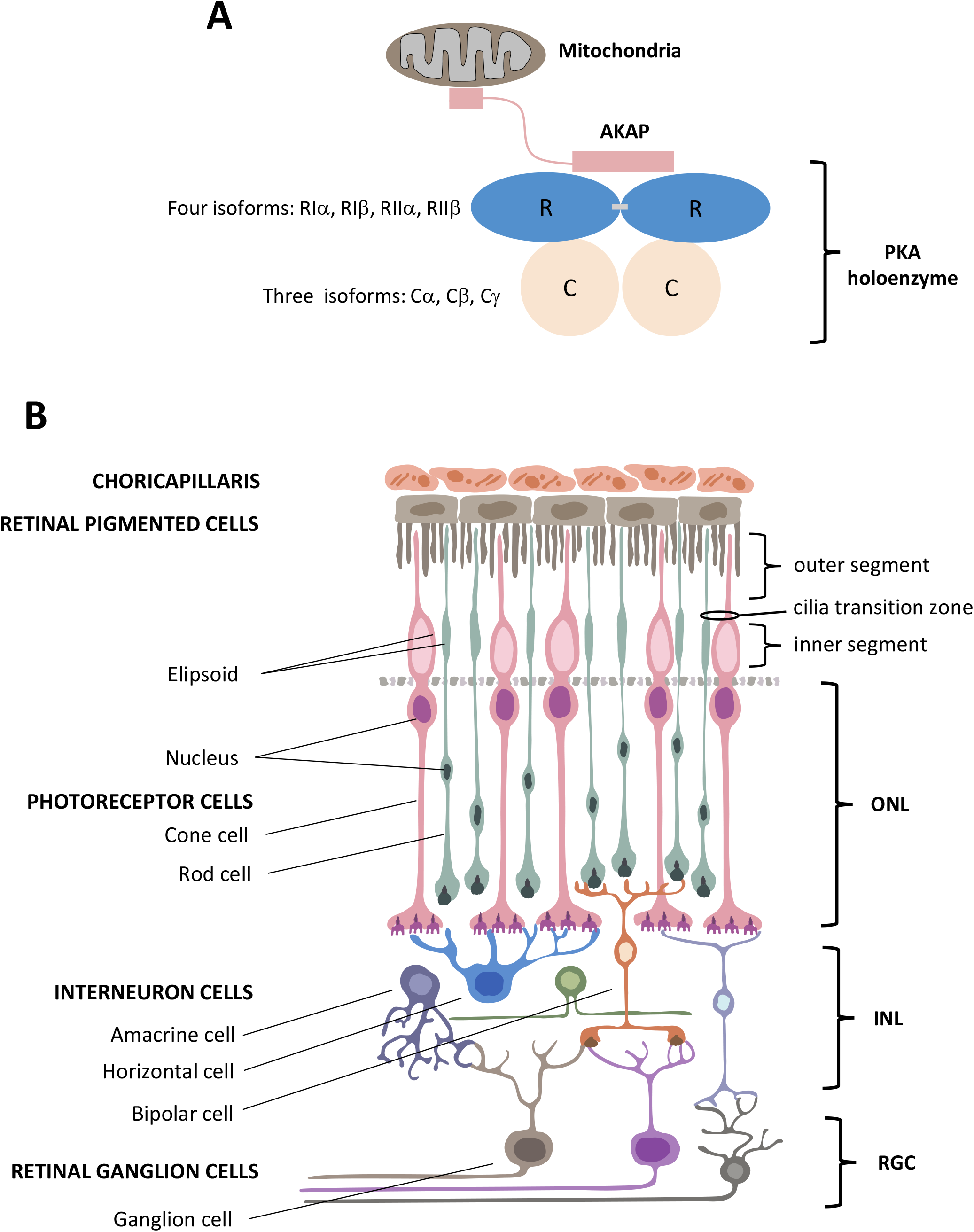
Structural organization of PKA and human retina. **A.** Schematic of PKA. In the inactive state two catalytic subunits are bound by a regulatory subunit dimer, which is typically anchored to intracellular locations such as the mitochondria by AKAP proteins. **B.** Retina consists of six neuronal cell types: cone, rod, horizontal, bipolar, amacrine, and ganglion cells. These six cell types are organized into three main layers within the retina that include the outer nuclear layer (ONL) with the nuclei of cones and rods, the inner nuclear layer (INL) with nuclei of interneuron horizontal, bipolar, and amacrine cells, and the retinal ganglion cell layer (RGC) that contains ganglion cell nuclei. Photoreceptor cone and rod cells directly accept photons in the specialized part of the cell called the outer segment, and they contain a mitochondrion-enriched ellipsoid in the inner segment, with the area connecting the outer and inner segment referred to as the cilia transition zone. RPE cells (top) are phagocytosis-competent cells indispensable for daily outer segment renewal. For interneurons: horizontal cells communicate with photoreceptors; bipolar cells communicate with both photoreceptor and retinal ganglion cells; and amacrine cells communicate with retinal ganglion cells. Finally, retinal ganglion cells convey the visual information to the brain.

The retina, a highly organized and accessible part of the central nervous system, is composed of three distinct cell layers with six easily distinguishable neuronal cell types (**Figure 1B**). In this study we thus used the retina as a model system to explore subcellular localization of PKA, which we hope will serve as a window into PKA signaling in the brain. Using specific antibodies raised against Cα, Cβ, RIIα, and RIIβ, as well as several other cell-type specific and organelle markers, we detailed the spatial localization of PKA C and RII subunits in tissue sections of human retina and found distinct localization patterns for each PKA subunit. Our results also show that Cα is the only subunit of the four that is enriched in the photoreceptor outer segment and cilia transition zone while Cβ is associated primarily with mitochondria. Cα, RIIα and RIIβ are specifically excluded from mitochondria. In addition, using novel RNA BaseScope technology, we achieved the first insights into the cell-type specific distribution of the Cβ splice variants in neurons.

## Results

### PKA C and RII subunits are differentially localized in human retina

To visually distinguish the different cell layers of the retina we used PKCα as a standard marker for rod bipolar cells. Our results show that the Cα subunit of PKA is expressed in the cell body of all cell types in the retina, (**Figure 2A**); however, in photoreceptors, it is enriched in the plasma membrane of the outer segment and in the cilia transition zone that connects the inner and outer segment. Notably, Cα was excluded from the membranes of the outer segment intracellular space, which contains photoreactive pigments (photoreceptor discs) (**Figure 2B, Sup Figure 1A-B**). Interestingly, Cα also co-localized with PKCα in rod bipolar cells.

**Figure 2.**
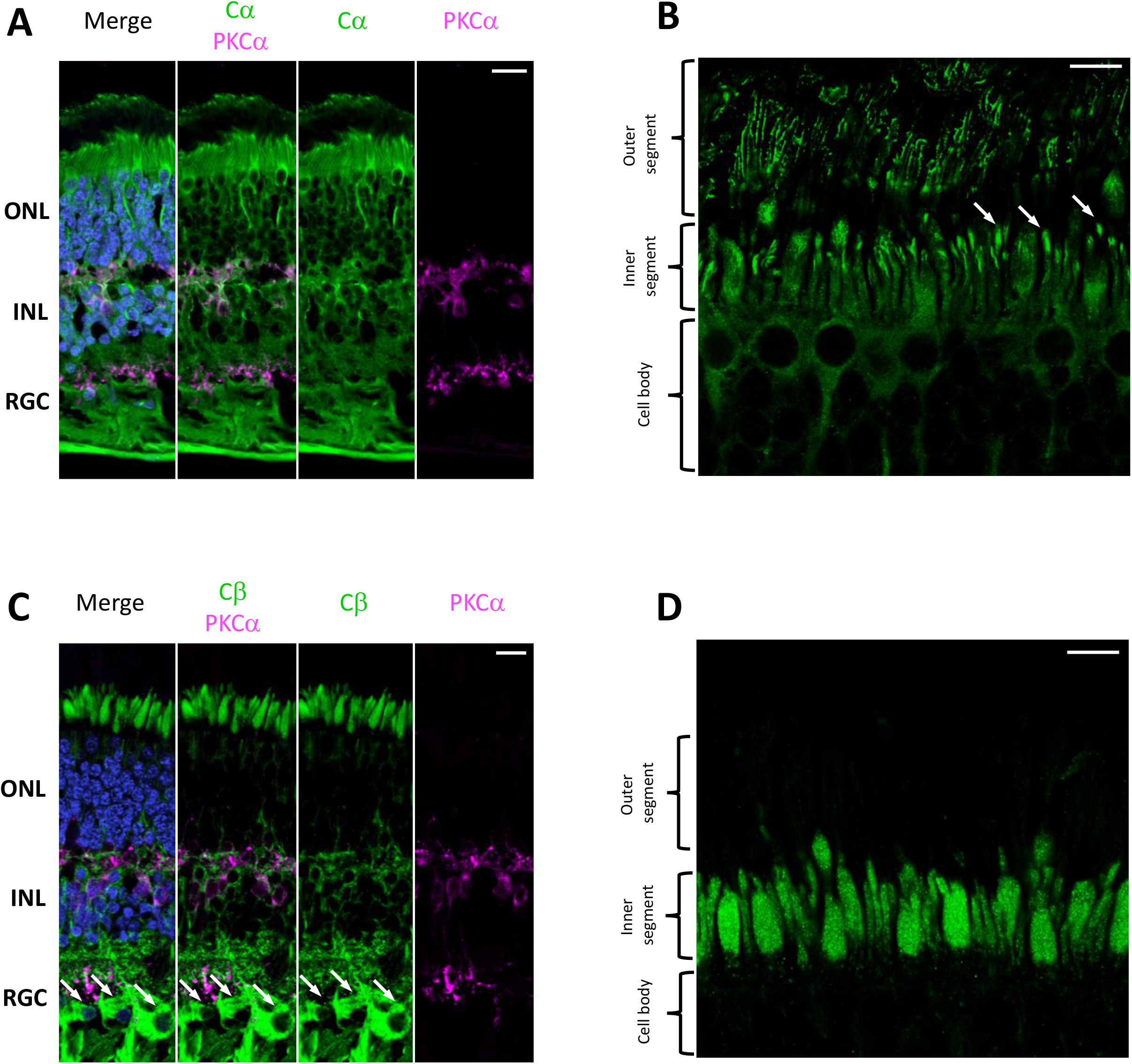
Cα and Cβ are differentially localized in human retina. **A-D**. All sections were labeled with anti-PKCα antibodies in order to identify rod-bipolar cells (purple), which helped distinguish between the outer nuclear layer (ONL), inner nuclear layer (INL), and retinal ganglion cell layer (RGC). Nuclei in blue, scale bar = 20μm (A, C), 10μm (B, D). **A**. Cα (green) is localized to the ONL, INL, and RGC, with strong signal in photoreceptor and PKCα-labeled rod bipolar cells (purple), and reduced signal in retinal ganglion cells **B.** In photoreceptors, Cα is generally localized to the cell body of both cone and rod photoreceptor cells; however, there is increased signal at the cilia transition zone (arrows) and outer segment membrane. Although arrows highlight a few examples, please note that all rods and cones contain this Cα localization in the cilia transition zone. **C.** Cβ (green) is localized to the ONL, INL, and RGC. with strong signal in photoreceptors, PKCα-labeled rod bipolar cells (purple), and retinal ganglion cells (arrows). **D.** In photoreceptors, Cβ (green) was limited to the ellipsoid of the inner segment, and notably absent from the outer segment and cilia transition zone.

Most striking in our analysis is that Cβ is localized in a manner that appears to be non-overlapping with Cα. In contrast to Cα, which is generally localized in the cell body of most cells in the retina, Cβ is enriched in photoreceptor and retinal ganglion cells (**Figure 2C**). Interestingly, the subcellular distribution of Cα and Cβ is also very different; unlike Cα, Cβ is not localized in the photoreceptor outer segment and is absent from the photoreceptor cilia transition zone. Instead, Cβ is localized almost exclusively to the inner segment of photoreceptors, most notably in the ellipsoid where there is a high concentration of mitochondria (**Figure 2D, Sup Figure 1C-D**). Cβ also seems to be present in axons and enriched in the synapses of photoreceptors where there are additional mitochondria. The striking differences in the subcellular localization of Cα and Cβ suggests strongly that these isoforms serve non-redundant functions. However, like Cα, Cβ is also partially co-localized with PKCα in rod bipolar cells.

With respect to regulatory subunit expression, RIIα and RIIβ are both localized to the cell body and axon of photoreceptor cells (**Figure 3A-D**); however, RIIβ is mostly detectable in cone cells and, in particular, in regions that surround the nucleus (**Figure 3D**). In contrast, while RIIα is also similarly enriched of cone cells, it is clearly expressed throughout the inner segment region of rod cells (**Figure 3B**). It is important to note that neither RIIα nor RIIβ are localized to the photoreceptor outer segment or to the cilia transition zone. The two RII subunits are also not highly expressed in retinal ganglion cells; and similar to Cα and Cβ, both RIIα and RIIβ are partially co-localized with PKCα in rod bipolar cells.

**Figure 3.**
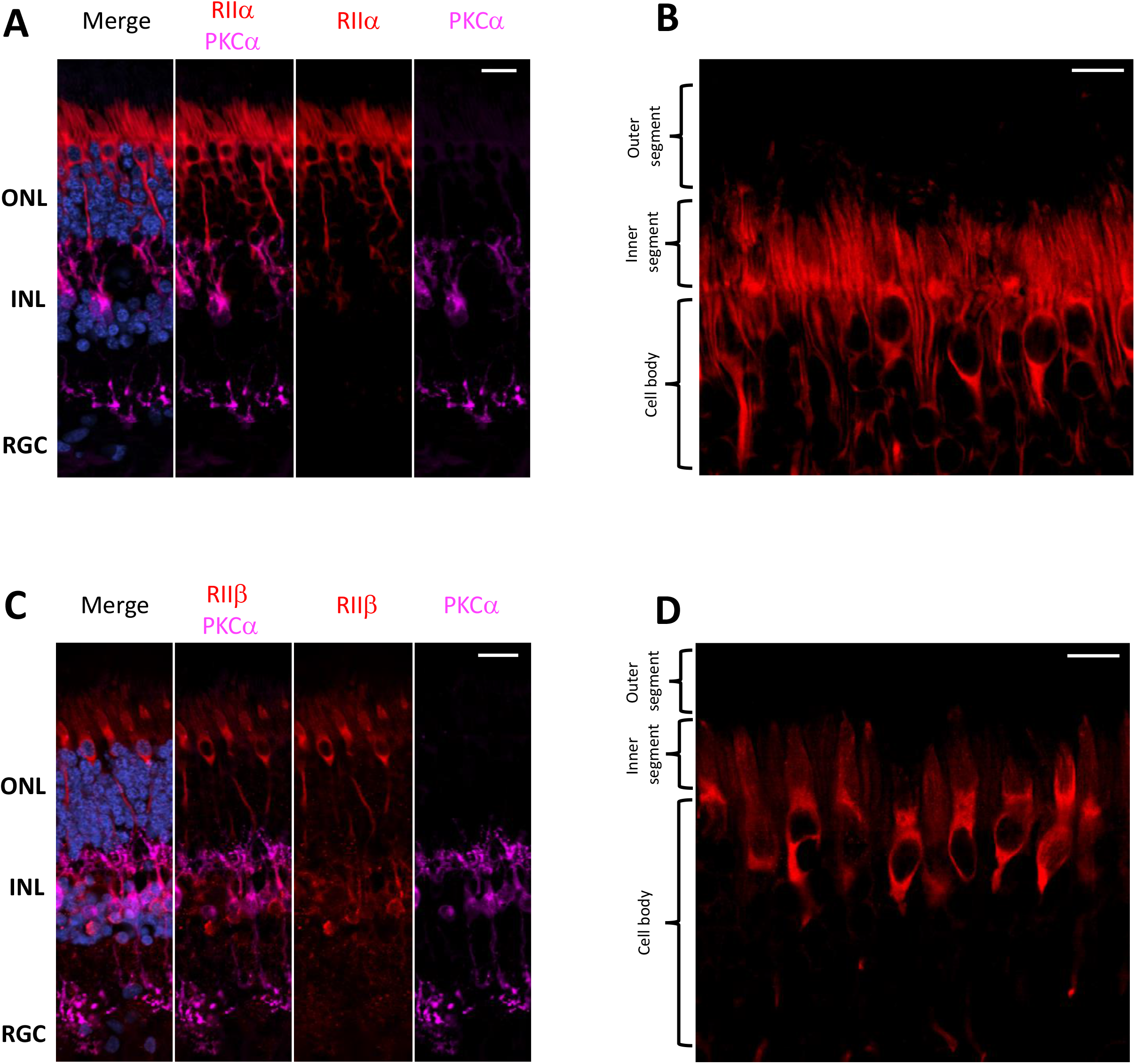
RIIα and RIIβ are differentially localized in human retina. **A-D.** All sections were labeled with anti-PKCα antibodies in order to identify rod-bipolar cells (purple), which helped distinguish between the outer nuclear layer (ONL), inner nuclear layer (INL), and retinal ganglion cell layer (RGC). Nuclei in blue, scale bar = 20μm (A, C), 10μm (B, D). **A**. RIIα (red) is localized to the ONL and INL, but not the RGC, with strong signal in photoreceptors and PKCα-labeled rod bipolar cells (purple). **B.** In photoreceptors, RIIα (red) is generally localized to the cell body and axons of both cone and rod photoreceptor cells. **C.** RIIβ (red) is localized to the ONL and INL, but not the RGC. There was strong signal throughout photoreceptor cell body and axons, as well as in interneuron cells where localization of RIIα was present in PKCα-labeled (purple) rod bipolar cells and non-PKCα-labeled interneuron cells. **D.** In photoreceptors, RIIβ (red) localization was distinct and strongest in the cell body and axons of cone photoreceptors.

### PKA C and RII subunits show distinct patterns of co-localization

Cα and RIIα are clearly co-localized in the cell body, axons, and synapses of photoreceptor cells, as well as in PKCα-expressing rod bipolar interneuron cells (**Figure 4A**) which would be consistent with these subunits existing as a holoenzyme complex. In photoreceptors, this co-localization is reduced with (as previously discussed) Cα being the only subunit localized to the cilia transition zone and outer segment (**Figure 4B, Sup Figure 2**). In contrast to Cα, Cβ is not co-localized with RIIα; although both seem to be expressed in photoreceptors and interneurons (**Figure 4C**). Subcellular localization further confirms these differences in photoreceptors, with localization of Cβ limited to the inner segment ellipsoid and RIIα distributed throughout the cell body and axon (**Figure 4D, Sup Figure 3**). RIIα is also absent from RGCs or at least expressed at very low levels whereas Cβ is enriched in most RGCs. Finally, RIIα and RIIβ are co-localized in photoreceptor and interneuron cells (**Figure 4E**); however, there is increased co-localization of both subunits in the cell body and axons of cone cells, and both RIIα and RIIβ protein expression appears to be somewhat polarized in cone cells with stronger localization near the nucleus around the ER/Golgi region of the cell (**Figure 4F, Sup Figure 4**).

**Figure 4.**
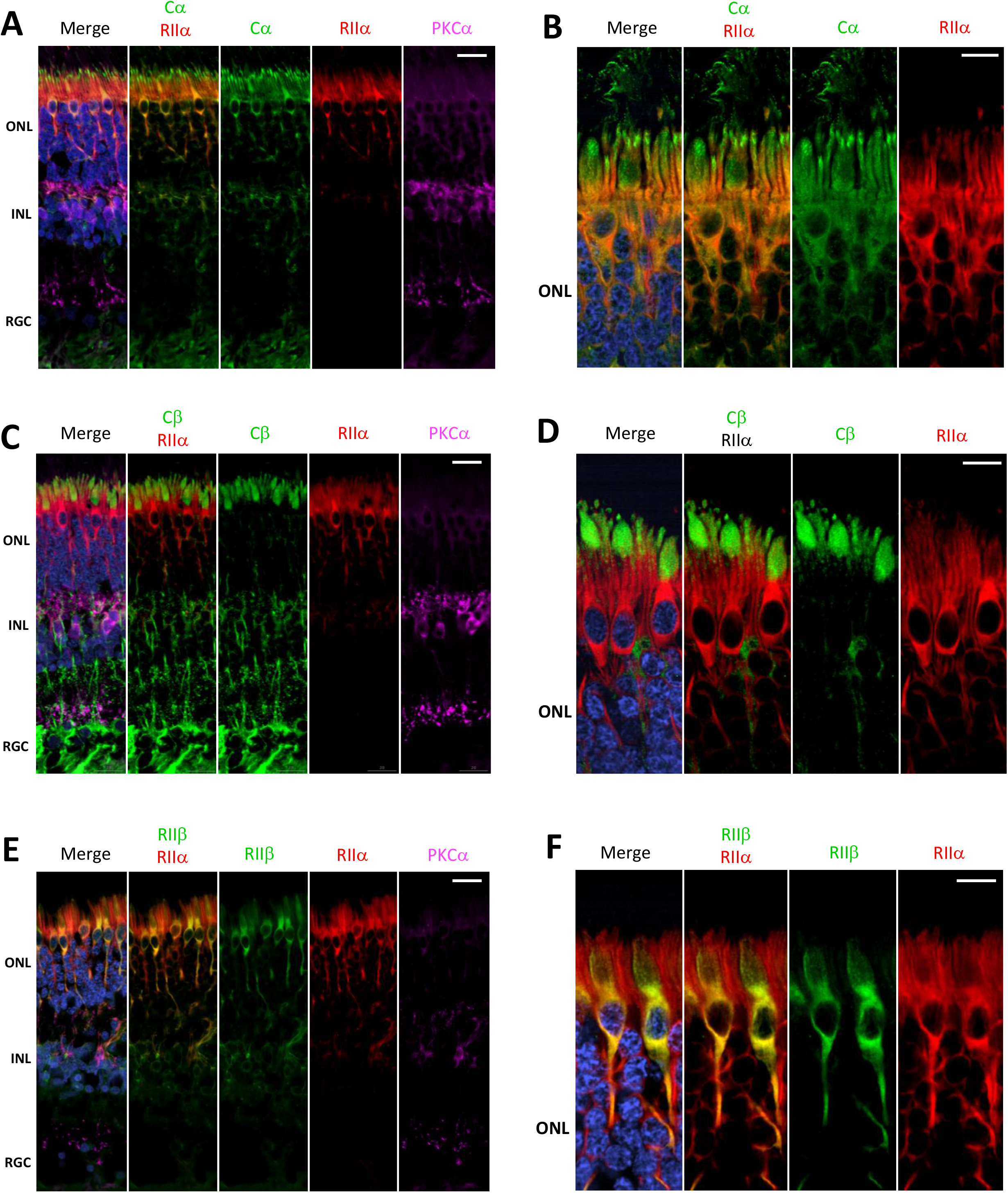
Cα, but not Cβ, co-localizes with RIIα; RIIα and RIIβ show similar localization. **A-F.** All sections were labeled with anti-PKCα antibodies in order to identify rod-bipolar cells (purple), which helped distinguish between the outer nuclear layer (ONL), inner nuclear layer (INL), and retinal ganglion cell layer (RGC). Nuclei in blue, scale bar = 20μm (A, C, E), 10μm (B, D, F). **A.** Cα (green) and RIIα (red) co-localized in cells in the ONL and INL, with strong signal overlap (orange, RIIα) in photoreceptors and in PKCα-labeled rod bipolar cells (purple). Only Cα was localized to the RGC. **B.** In photoreceptors, PKA-Cα (green) and PKA-RIIα (red) are clearly co-localized in the cell body and axons (orange, Cα RIIα); however, only Cα was present in the cilia transition zone and outer segment membrane. **C.** Cβ (green) and RIIα (red) intracellular localization differs in cells in the ONL and INL, with no signal overlap (Cβ RIIα) in photoreceptors nor in PKCα-labeled rod bipolar cells (purple). Only Cβ was localized to the RGC. **D.** Cβ (green) and RIIα (red) are both clearly localized in photoreceptors; but Cβ (green) is found in the inner segment, while RIIα (red) is found in the cell body and axons, with no signal overlap (Cβ RIIα). **E.** RIIβ (green) and RIIα (red) co-localized in cells in the ONL and INL, with strong signal overlap (orange, RIIβ RIIα) in photoreceptors and in PKCα-labeled rod bipolar cells (purple); and neither RIIβ and RIIα were localized to the RGC. **F.** In photoreceptors, RIIβ (green) and RIIα (red) clearly co-localized to the cell body and axons (orange, RIIβ RIIα); however, RIIβ was distinctly enriched in cone photoreceptor cells, while RIIα was diffusely distributed in both rod and cone photoreceptor cells.

### Cβ is localized with mitochondria

Given that Cβ is so highly expressed in the inner segment ellipsoid we wanted to explore co-localization of the PKA catalytic subunits with mitochondria. For this we used anti-OxPhos Complex V subunit I antibodies created against mitochondrial encoded cytochrome C oxidase 1 (MT-CO1) that binds to the outer mitochondrial membrane. Our results show that Cα is not localized to mitochondria in photoreceptors, interneurons, and retinal ganglion cells (**Figure 5A, Sup Figure 5**). Specifically in photoreceptors we found Cα was notably absent from the mitochondria-enriched ellipsoid of the inner segment (**Figure 5B**), which was confirmed in a single compiled z-stack cross section as well as in nine individual z-stack images (**Figure 5C, Sup Figure 6**).

**Figure 5.**
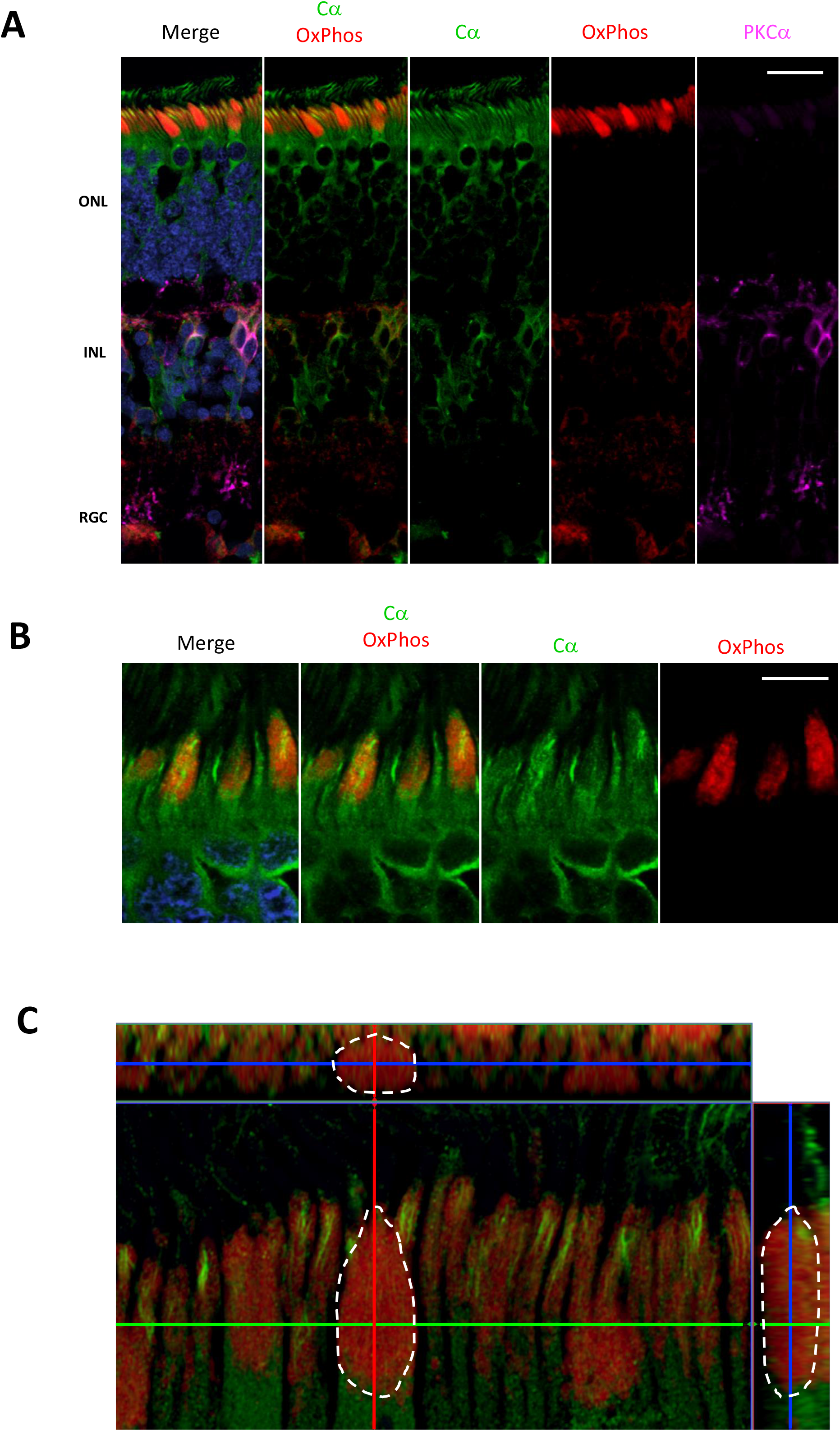
Cα localization is distinctly separate from mitochondria. **A-C.** All sections were labeled with anti-PKCα antibodies in order to identify rod-bipolar cells (purple), which helped distinguish between the outer nuclear layer (ONL), inner nuclear layer (INL), and retinal ganglion cell layer (RGC). Nuclei in blue, scale bar = 20μm (A), 10μm (B). **A.** Cα (green) was not localized with OxPhos(red)-labeled mitochondria in the ONL, INL, and RGC. **B.** In photoreceptors, Cα (green) was distinctly separate from OxPhos (red)-labeled mitochondria. Cα signal was strongest in the cell body, axon, and cilia transition zone, while OxPhos signal was enriched in the mitochondria-containing ellipsoid of the photoreceptor inner segment. **C.** Cα (green) is notably absent, while OxPhos(red)-labeled mitochondria fill the entire inner segment ellipsoid of a single cone cell (outlined).

On the other hand, Cβ was localized almost exclusively to mitochondria, including co-localization in photoreceptor, interneuron, and retinal ganglion cells (**Figure 6A, Sup Figure 7**). The signal from anti-Cβ antibodies completely overlapped with mitochondria identified using anti-OxPhos antibodies in the photoreceptor inner segment (**Figure 6B**). This was further confirmed using multiple z-stack images that show co-localization in the ellipsoid was complete, which can be visualized in a single compiled z-stack cross section as well as throughout nine individual z-stack images (**Figure 6C, Sup Figure 8**). We also noticed OxPhos-positive small mitochondria below the primary mitochondria ellipsoid, which co-localize with Cβ, while additional small Cβ-positive puncta do not show this co-localization (**Sup Figure 7A**).

**Figure 6.**
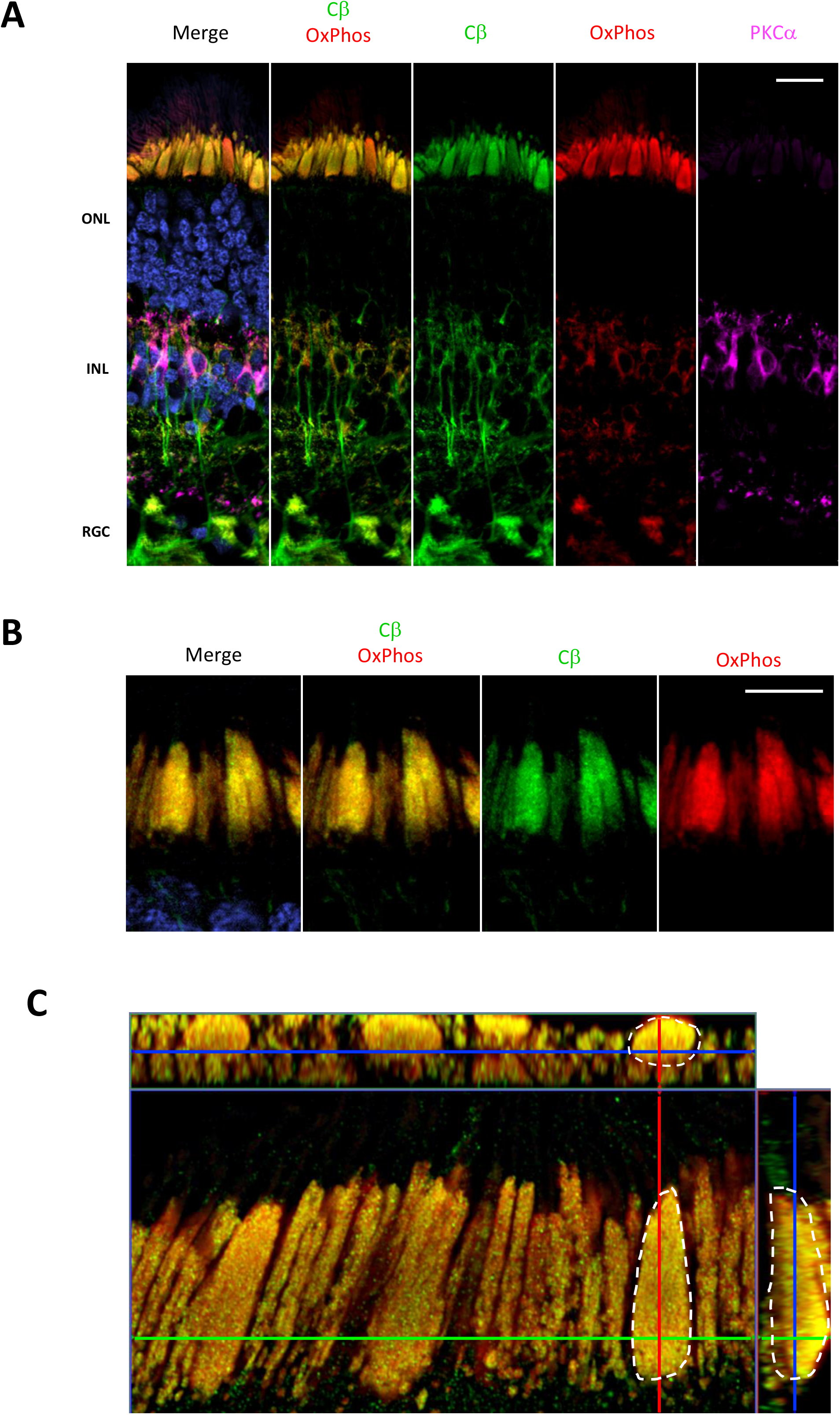
Cβ is co-localized with mitochondria. **A-C.** All sections were labeled with anti-PKCα antibodies in order to identify rod-bipolar cells (purple), which helped distinguish between the outer nuclear layer (ONL), inner nuclear layer (INL), and retinal ganglion cell layer (RGC). Nuclei in blue, scale bar = 20μm (A), 10μm (B). **A.** Cβ (green) and OxPhos(red)-labeled mitochondria co-localize in the ONL, INL, and RGC, with clear signal overlap (yellow, Cα OxPhos) in every cell layer. **B.** In photoreceptors, Cβ (green) and OxPhos(red)-labeled mitochondria are distinctly co-localized to the inner segment ellipsoid. Cβ is limited to this region, and does not localize to cell body, axon, and cilia transition zone. **C.** Cβ and OxPhos-labeled mitochondria co-localize (yellow) to fill the entire inner segment ellipsoid of a single cone cell (outlined).

### Cβ splice variants are differentially expressed across the retina

Western blots of human retina samples show two bands for Cβ, while only one band is present for Cα (**Figure 7A**). These results highlight the presence of multiple Cβ isoforms in human retina, a finding that is further supported by a previous study using RNA hybridization that revealed tissue-specific expression of multiple Cβ isoforms (Orstavik et al., 2001). As seen on the protein sequence alignments of Cα and the multiple Cβ isoforms, the variants in Cβ are due exclusively to differences in N-terminus (**Figure 7B**). Since they all have approximately the same molecular weight, they cannot be distinguished by gel migration nor with the current available antibodies that recognize an antigen common to all Cβ isoforms (Asp324). Because of the latter, immunohistochemistry cannot distinguish between different Cβ isoforms.

**Figure 7.**
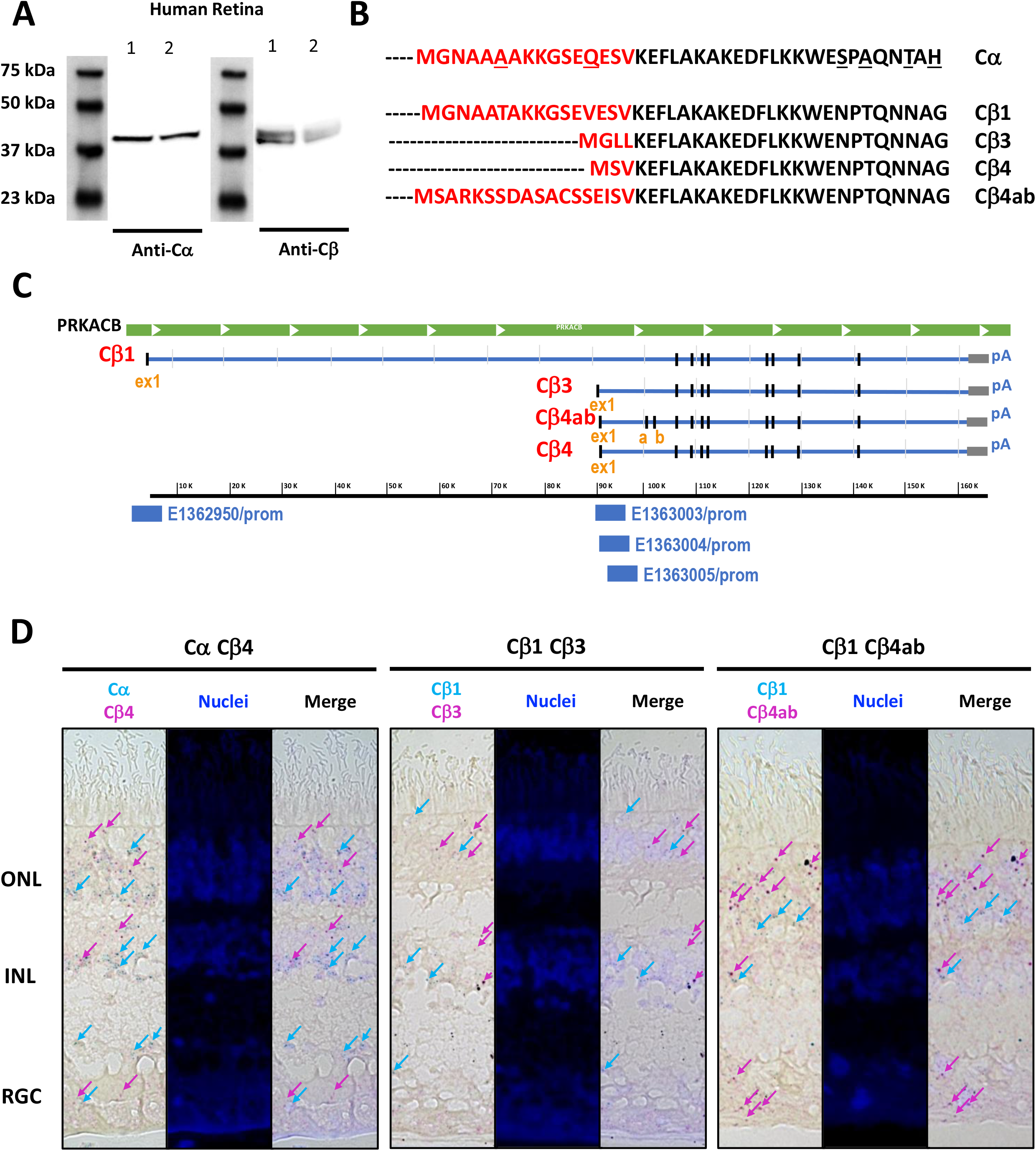
Similar Cα and Cβ proteins are present but multiple Cβ variants show variable expression across the human retina. **A.** Anti-Cα antibodies recognized a ~39 kDa band, and anti-Cβ antibodies recognized two bands at ~39 kDa and ~42 kDa bands, in Western blots from two samples (1, 2) of human retina tissue extracts. **B.** Protein sequence alignment of Cα and Cβ splice variants Cβ1, Cβ3, Cβ4, and Cβ4ab, and **C.** Corresponding RNA sequence alignments of Cβ isoforms, with unique promoter sequences identified for each variant explored. ex1 – exon 1; a, b – exon a, exon b. **D.** BaseScope Duplex assay results show Cα, Cβ1, Cβ3, Cβ4, and Cβ4ab expression in photoreceptors in the outer nuclear layer (ONL) and interneurons in the inner nuclear layer (INL). Cβ4 and Cβ4ab were also expressed in the retinal ganglion cell layer (RGC). Expression of Cα and Cβ1 highlighted by blue arrows, and expression of Cβ3, Cβ4, and Cβ4ab highlighted by magenta arrows.

**Figure 8.**
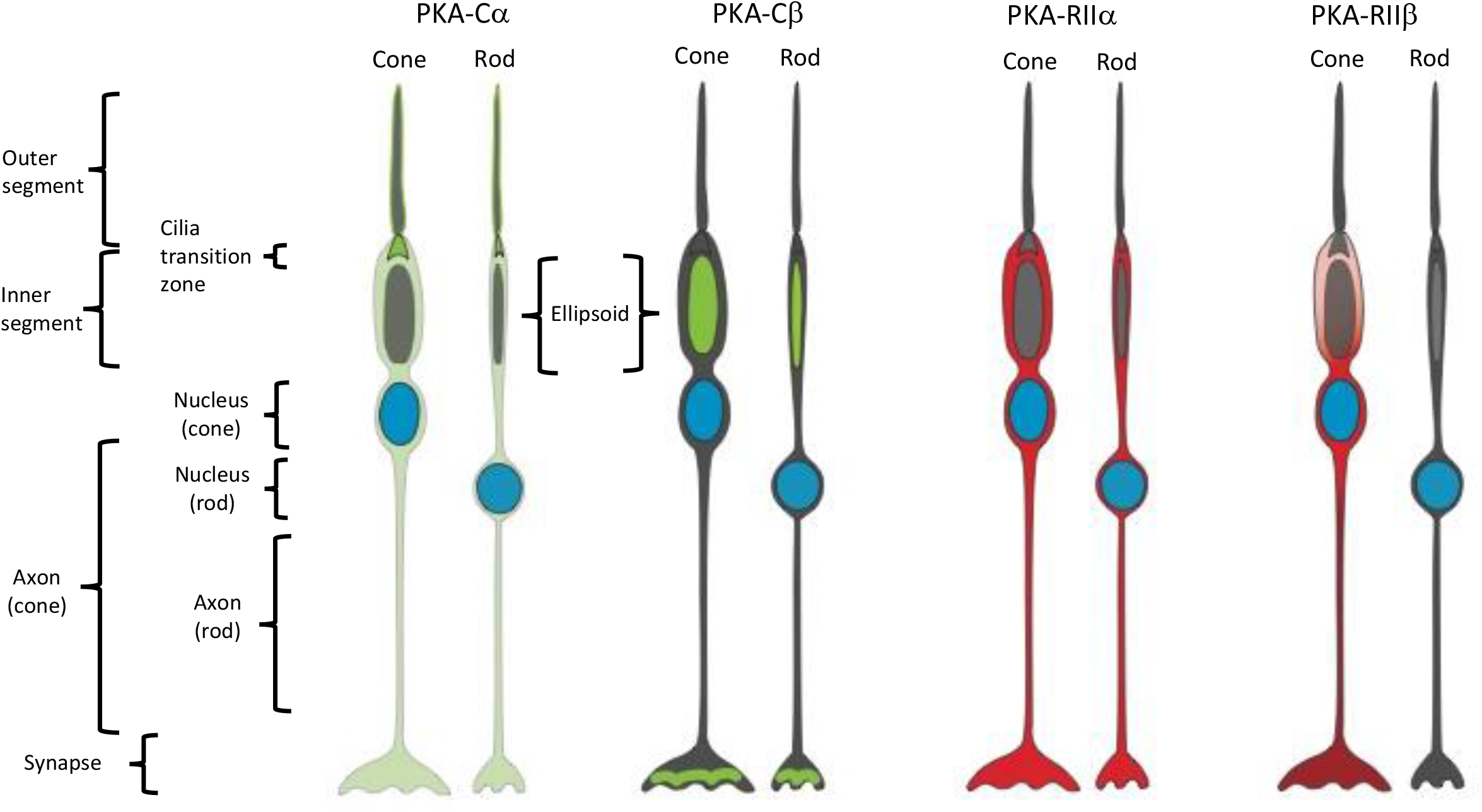
Summary of PKA expression in cone and rod photoreceptor cells. Cα: outer segment membrane, cilia transition zone, cell body, axon, and synapse of cone and rod cells. Cβ: inner segment mitochondrion-enriched ellipsoid of cone and rod cells, as well as in the synaptic region. RIIα: cell body, axon, and synapse of cone and rod cells. RIIβ: cell body, axon, and synapse of cone cells.

When examining the Cβ locus we noticed that each variant has a unique exon one, further confirmed by the ENCODE *in silico* prediction of promoter sequences (ENCODE Consortium Project et al., 2020) coinciding with each first exon, and with additional splice variants that shuffle short exons (4 vs. 4ab) (**Figure 7C**). RNA sequence alignments detected differences in Cβ1, Cβ3, Cβ4, and Cβ4ab at the N-terminus that were sufficient to design BaseScope probes specific for each isoform (**Sup Figure 9**). Our *in-situ* hybridization results show that Cα is expressed in all retinal cell layers, which is consistent with previous knowledge about it being ubiquitously expressed in the body. Interestingly, Cβ isoforms displayed more distinctive expression pattern. While Cβ4 and Cβ4ab are expressed in all retinal cell types, Cβ1 seems to be less abundant and Cβ3 is practically excluded from RGCs (**Figure 7D, Sup Figure 10**).

## Discussion

Using the retina with its highly differentiated and spatially organized neurons as an experimentally accessible “window into the brain”, we studied the expression of PKA catalytic (C) and regulatory (R) subunits where Cβ, in particular, is predicted to be highly expressed but previously unexplored. In our study, we not only found that Cβ is indeed highly expressed in these neurons, but also that its subcellular localization is distinct from that of Cα. Most notable is the exclusive localization of Cβ to mitochondria. While Cα is widely expressed in a diffuse fashion in the cell bodies, it is specifically excluded from mitochondria. In turn, Cα is the only subunit enriched in the plasma membrane of the photoreceptor outer segment and in the cilia transition zone. With regard to the regulatory subunits, we explored RIIα and RIIβ and also found different localization patterns for each. Like Cα, RIIα and RIIβ are expressed in cell bodies but are excluded from mitochondria; however, unlike Cα, they are also excluded from the photoreceptor outer segment and cilia transition zone. We also found that RIIα was evenly distributed throughout the cell body of cone and rod cells, while RIIβ was enriched in cone cells. Cβ also does not appear to be co-localized with either RIIα or RIIβ. Finally, using RNA BaseScope technology to discriminate between the multiple Cβ isoforms, we found differences in their spatial and cellular distribution, which suggests that the Cβ isoforms may also represent functionally non-redundant family of proteins.

### Cβ in mitochondria

Both Cα and Cβ subunits seem to be expressed in all neurons; however, we show striking co-localization of Cβ with mitochondria, while Cα appears to be excluded from this organelle. Localization of Cβ to mitochondria was confirmed by demonstrating co-localization with cytochrome c oxidase, the terminal complex of the electron transfer chain. Mitochondrial localization of Cβ is most noticeable in the large ellipsoid body of cone cells due to the presence of densely packed mitochondria, while in other cells such as the retinal ganglia cells, the mitochondria and Cβ are still co-localized but are distributed throughout the cytoplasm. Additional enrichment of Cβ was detected in mitochondria localized to photoreceptor synapses and RGC axons.

It has been previously shown that PKA is an essential regulator of different aspects of mitochondrial biology (Feliciello et al., 2005). Multiple lines of evidence show the pivotal role of PKA in mitochondrial morphology, dynamics, and turnover by phosphorylating dynamin-related protein 1 (Drp1) to inhibit mitochondrial fission (Edwards et al., 2020; Strack et al., 2013), the proapoptotic protein BAD to promote survival (Virdee et al., 2000), and mitochondrial-anchored cytoskeletal proteins to remodel dendrites (Dagda and Banerjee, 2015). Our data suggest that these functions are most likely delivered by the Cβ subunit of PKA, pinpointing the non-redundant roles of both Cα and Cβ. PKA has also been shown to have a potential role in the mitochondrial matrix, including phosphorylating a subunit of complex I (NDUFS4) *in vitro* and *in vivo* to elevate oxidative/phosphorylation (De Rasmo et al., 2008). Further studies are required to understand which Cβ isoform variant is involved in particular mitochondrial processes and where in the mitochondria the Cβ subunit is localized. Our imaging shows that Cβ is densely packed in the ellipsoid of cone cells but we do not distinguish whether localization is at the outer membrane, inner membrane, or inner membrane space. Going forward it will also be important to investigate the role of Cβ in human disease with specific regards to PKA signaling in mitochondria, which has been previously linked to several neurodegenerative diseases including Alzheimer’s, Huntington’s, and Parkinson’s Disease (Dagda and Banerjee, 2015).

### Cβ isoforms

Given that the Cβ transcriptional variants are so similar in sequence, differing only at the region corresponding to the first exon, it is not possible to discriminate between the Cβ family members with currently available antibodies. As a first step to decipher whether the Cβ protein variants are expressed in a cell-specific manner and whether they represent a set of functionally non-redundant proteins, we used RNA BaseScope technology, which allows for the detection of small differences in the RNA transcript. This novel approach allowed us to demonstrate that the Cβ variants are expressed in a cell-specific manner. In particular, while Cβ1 and Cβ3 are expressed in photoreceptors and interneurons, Cβ1 is particularly enriched in these cells. In contrast, while the Cβ4 and Cβ4ab are expressed in most retinal cells, they appear to be the primary Cβ isoforms expressed in RGCs. This suggests that specific Cβ protein variants have cell-type specific roles and that their distribution may indicate particular cellular functions. It is also important to note here that while most PKA disease mutations are associated with either Cα (Cushing’s Disease) or RIα (Carney Complex Disease or Acryodysostosis), a recent report described patients with four different Cβ mutations who all showed a severe and complex phenotype that includes skeletal and cardiac defects as well as a polydactyl presentation (Palencia-Campos et al., 2020). In fact, these mutant proteins inhibited Sonic hedgehog signaling, which supports a role for Cβ isoforms in neurodevelopment.

### Cα is enriched in the plasma membrane of the outer segment and in the cilia transition zone of photoreceptor cells

Interestingly, of all the subunits explored in this study, Cα was the only one found to be highly enriched in the photoreceptor outer segment plasma membrane and in the cilia transition zone, also known as the connecting cilium. With regard to the outer segment membrane, this sequestration of a PKA C subunit to the cell membrane could be related to the recent report showing that the PKA C subunit is directly bound in a non-canonical way to Smoothened, a GPCR that is linked to the Sonic hedgehog pathway and is localized to cilia (Arveseth et al., 2021). Additionally, Cα localization is especially prominent at the tip of the outer segment membrane where photoreceptor cells interact with the retinal pigment epithelium (RPE) cells. RPE cells are responsible for removal of outer segment material and degraded cell components daily, and with the supposed high mitochondrial turnover rate in photoreceptor cells it is possible that the PKA pathway could play a role in photoreceptor regeneration via RPE cells. The particular enrichment of Cα in the cilia transition zone is interesting in the context of mutations of several proteins associated with Usher syndrome previously indicated to be involved with the PKA pathway (Kremer et al., 2006, Mathur and Yang 2015) and provides novel opportunities to study the mechanism of this disease which is a common cause of deafness and blindness.

### RIIα vs. RIIβ

With respect to R subunits, this study focused on the RII class of regulatory subunits that include RIIα and RIIβ. Unlike the pseudo-substrate nature of the RI subunits, RII subunits both inhibit activity of and are phosphorylated by the C subunits (Taylor et al., 2012). In addition, the RII subunits are mostly targeted through AKAPs to receptors, ion channels, and transporters placing them in close proximity to their target substrates. Early small angle X-ray scattering studies (Vigil et al., 2004, 2006) and the recent cryoEM structure of the RIIβ holoenzyme show that these two holoenzymes are structurally quite distinct (Lu et al., 2020). Our results further support the functional non-redundancy of RIIα and RIIβ as we show distinct localization patterns for each.

Specifically in photoreceptors where RIIα localization was readily detectable in rods and cones but RIIβ localization enriched in cones, which might suggest that RIIα provides some additional light/dark related regulation in rod cells. Interestingly, both RIIα and RIIβ localization in cones displayed distinct enrichment around the nucleus where Golgi and ER are typically enriched. It is important to note that a significant limitation to our study is that we could not explore RIα and RIβ in our human retina samples as our antibodies were unable to determine their subcellular localization. We do not presume that neither subunit is present in retinal cells, but we do suggest these subunits might be better investigated in traditional model species where tissue fixation and pretreatment methods are more easily modified, unlike treatments of human samples that are understandably less flexible due to the nature of tissue availability.

### Conclusion and future perspectives

While localization of functionally non-redundant PKA holoenzymes and R subunits by AKAPs has been widely recognized as a way of achieving specificity in PKA signaling, the role of C subunit specificity has been largely ignored. Although early hybridization studies showed decades ago that over half of PKA signaling in the brain is mediated by Cβ, little attention was paid to the specific localization and functional roles of the Cβ proteins which include several sub families as well as multiple splice variants. Our demonstration of distinct patterns of localization in the highly specialized neurons of the retina suggests that this is a rich world of PKA signaling that has yet to be explored; and our results highlight that it is essential to explore localization of endogenous proteins in tissues as a companion to structural analysis of signaling proteins like PKA and traditional exploration of protein function using various immortal cell lines. It is also important to consider tissue-specific localization of isoforms in conjunction with mass spectrometry analyses; both are essential. The expanded combinatorial diversity of PKA signaling that is introduced by the diversity of the C isoforms exemplifies why these approaches need to be rigorously carried out in parallel. Based on our imaging, Cα and Cβ are clearly functionally non-redundant; while our preliminary mapping of the expression of different Cβ transcripts with BaseScope strongly suggests that the Cβ variants may also be functionally non-redundant. Clearly an important next step is to determine where the Cβ subunit is localized in the mitochondria and what specific Cβ variants are there; and some of our future challenges include understanding how they contribute specifically to cell function and to mitochondrial homeostasis, and to what extent they play a role in the recently discovered Cβ disease phenotypes.

## Materials and methods

### Antibodies

Primary antibodies were generally used at 1:100 dilution, including: custom rabbit serum anti-Cα antibodies; rabbit polyclonal anti-Cβ antibodies (Lifespan Biosciences, catalog # LS-C191947); mouse monoclonal anti-RIIα antibodies (Santa Cruz Biotechnology, catalog # sc-137220, RRID: AB_2268608); rabbit monoclonal anti-RIIβ antibodies (Abcam, catalog # AB75993, RRID: AB_1524201); and AF647 conjugated anti-PKCα antibodies (Santa Cruz Biotechnology, catalog # sc-8393, RRID: AB_628142). Custom mouse monoclonal anti-RHO (1D4 anti-rhodopsin) antibodies were used at 1:1000, and mouse monoclonal anti-OxPhos antibodies (Complex IV, Subunit I, Thermo Fisher Scientific catalog # 459600, RRID: AB_2532240) were used at 1:500. Peanut Agglutinin conjugated with rhodamine was used at 1:1000 to identify cones (Vector Laboratories, catalog # RL-1072-5). Secondary antibodies: Alexa Fluor 488 Donkey anti-rabbit (Jackson ImmunoResearch Laboratories, RRID: AB_2313584) and Alexa Flour 594 Donkey anti-mouse (Jackson ImmunoResearch Laboratories, RRID: AB_2338871) were used at 1:250. Nuclei were stained with DAPI (2μg/ml). Custom anti-Cα antibodies specifically recognized purified Cα protein at ~39 kDa but not purified Cβ protein, while pan anti-Cβ recognized purified Cβ protein at ~39 kDa but not purified Cα protein (**Sup Figure 11A**); and sections incubated with secondary antibodies as negative controls showed no signal (**Sup Figure 11B**).

### Patient Information

Human retina tissue sections and retinas extracts were obtained from a normal (age 81, 83 and 91 years) donors (San Diego Eye Bank, CA, USA) with appropriate consent from the San Diego Eye bank and with a protocol approved by the University of California, San Diego Human Research Protection Program. Donors have no history of eye disease, diabetes, or chronic central nervous system disease.

### Tissue Processing

After enucleation, eyeballs were fixed for ~24h in the 10% formalin. Next, the anterior segment, crystalline lens and vitreous were removed, and the eye cups were processed for cryostat sections (12 μm). Sections were stored in −80 until the use for experiment.

Frozen sections were defrosted (10 min, RT) and placed in 1X PBS (10 min, RT). Rehydrated sections were then blocked with 2% normal donkey serum, 0.02% keyhole limpet hemocyanin in 1X PBS-TX (0.2% Triton-X) for 1h. Sections were then incubated in primary antibodies overnight at 4 °C. Slides were then washed in 1X PBS (3x, 15min, RT) and sections were incubated in DAPI and secondary antibodies (2h, RT). Slides were then washed in 1X PBS (3x, 15 min, RT) and sections were permanently mounted in Fluorogel with tris buffer (Electron Microscopy Sciences, Hatfield, PA, USA).

### Imaging

High resolution images were captured using a laser scanning confocal mode (A1R HD, Nikon) on an Eclipse Ti2-E (Nikon) housed in the UCSD Nikon Imaging Center. Samples were excited with 405 nm, 488 nm, 561 nm, and 640 nm laser from a laser unit (LU-NV, Nikon) and emission captured using a slowed galvano scanner mode. Images were either acquired at 40x (S Fluor 40x NA 1.30 oil) or 100x (Plan Apo lambda 100x NA 1.45 oil). 40x images were used to determine general protein expression patterns, and 100x images were used to determine cell- and organelle-specific protein localization. Cα, Cβ, RIIα, RIIβ, OxPhos, RHO, and PNA image acquisition details: Laser Power 3.0%, High Voltage 10-25, Offset 5, LUT range 100-3000. PKCα image acquisition details: Laser Power 4.5 %, High Voltage 90-100, Offset 0, LUT range 50-2000. DAPI image acquisition details: Laser Power 5.0%, High Voltage 60, Offset 5, LUT range 60-745. Pixel size was set to 17 nm, and pixel dwell time was >1 μs with unidirectional scan mode. Digital images were adjusted for brightness and contrast only, using Omero Insight software (University of Dundee & Open Microscopy Environment). Antibody specificity was verified on control sections incubated with only secondary antibodies and similarly processed, including image acquisition parameters and post-processing in Omero Insight.

To obtain 3D images of the tissue, additional high-resolution images were taken using a ZEISS LSM880 with Airyscan at the La Jolla Institute for Immunology. Samples were excited with 405, 488, and 561 nm laser light with main beam splitters set to 405 nm, and 488/561/633 nm. Laser power was kept below 5%. Images were acquired with Plan-Apochromat 40x NA 1.4, Plan-Apochromat 63x NA 1.4, or Alpha Plan-Apo 63x NA 1.46 oil objectives using Immersol 518F 30°C immersion oil. For imaging DAPI, emitted fluorescence was filtered by a bandpass 420-480 nm + longpass 605 nm filter. Detection of Alexa Fluor 488 utilized a bandpass 420-480 + 495-550 nm filter, and fluorescence of Alexa Fluor 568 was collected with 570 nm longpass filter and a bandpass 420-480 + 495-620 nm filter. Pixel size was set to 40 nm, and pixel dwell time was >1 μs with unidirectional scan mode. Z-stacks were acquired with optimal step size according to ZEN Black 2.3 SP1 software. Airyscan detector was automatically aligned and run in the superresolution mode with gain settings between 700 and 750V to achieve optimal dynamic range. Data were processed with automatic Airyscan processing settings and resulting 16-bit images were adjusted for brightness and contrast using ZEN Blue 3.1 lite software. Maximum intensity or orthogonal projections were created for selected images.

### Western blots

Retina tissue were lysed with RIPA buffer to get the whole cell lysate. 50 μg of each sample was separated on a 10% gel and blotted for probing with anti-Cα and Cβ, respectively. 30ng of purified Cα and Cβ was also included in the gel as specific control.

### BaseScope duplex assay

BaseScope duplex assay for the mRNA detection was performed following the manual of BaseScope duplex detection kits (ACD, 323870) to detect splicing variants of PKA catalytic subunit beta within which, four of them bare a difference with only a few nucleotides. Fresh frozen sections were rinsed in PBS and baked at 60°C for 1 hour followed by dehydration in 50%, 75% and 100% ethanol. Formaldehyde-fixed paraffin-embedded (FFPE) sections were 60°C baked and deparaffinized twice in xylene and twice in ethanol. Sections were dried and treated with hydrogen peroxide for 10 min, then the samples were heated in targets retrieval buffer at 99°C for 15 min followed by incubation for 3 min in ethanol. Protease treatment was performed using protease III at 40 °C for 30min. Sections were washed in water and the target mRNA were hybridized with probes by incubation at 40 °C for 2 h. Probes were designed and produced by Advanced Cell Diagnostics. Sections were washed in wash buffer followed by a cascade of hybridization steps with signal amplification molecules and added green and red substrates to the targets. Both the steps for Amp7 and Amp11 in BaseScope duplex detection kits manual were prolonged to 45 h. After all the hybridization steps, nuclei in the sections were stained with DAPI, dehydrated with xylene, and mounted with VectaMount medium (Vector Labs).

Probes used in the assay:

**Cα1** (NM_002730.4): BA-Hs-PRKACA-3zz-st, Cat#868751, Lot#20164A

**Cβ1** (NM_002731.3): BA-Hs-PRKACB-tv2-2zz-st, Cat# 867501, Lot#20164A

**Cβ3** (NM_001242858.2): BA-Hs-PRKACB-tv5-2zz-st-C2, Cat#868741, Lot#20164A

**Cβ4** (NM_001375565.1): BA-Hs-PRKACB-tv18-1zz-st-C2, Cat#867391-C2, Lot#20164A **Cβ4ab** (NM_001375560.1): BA-Hs-PRKACB-tv13-1zz-st-C2, Cat#868731-C2, Lot#20164A

## Acknowledgements

JNR was supported by NIH/NIGMS IRACDA K12 GM068524. Work in the DSK laboratory was supported by R01 EY027011, RPB Special Scholar Award, Glaucoma Research Foundation Shaffer Grant and in part by an unrestricted grant from Research to Prevent Blindness (New York, NY) awarded to Department of Ophthalmology, UC Irvine. Work in the SST laboratory was supported by 1R35 GM130389 and R03 TR002947. The Zeiss LSM 880 was funded by NIH S10OD021831. We would like to thank Dr. Eric Griffis (UC San Diego) and the Nikon Imaging Center at UCSD for the assistance with imaging, Eva Henry-Dawson for her schematic drawing of retina and photoreceptors, and Dr. Benjamin Myers (University of Utah) for providing critical feedback on the manuscript draft.

## Competing interests

The authors declare no competing interest.

## Author contributions

SST and DSK conceived the project, guided experimental design, and interpretation of the data. RI began to explore retina as a model system; and RI and YM preformed original exploratory imaging. JNR conducted all IHC experiments for the manuscript and most of the imaging; ZM conducted the Airyscan imaging. YM conducted the western blots experiments and prepared images for Figure 7A. YM, QX, and JNR conducted BaseScope experiments and prepared the BaseScope figures. JNR prepared all the figures for the manuscript. JNR, SST, and DSK wrote the manuscript. All authors read and edited the manuscript.

## Supplemental Figure Legends

**Supplemental Figure 1.**
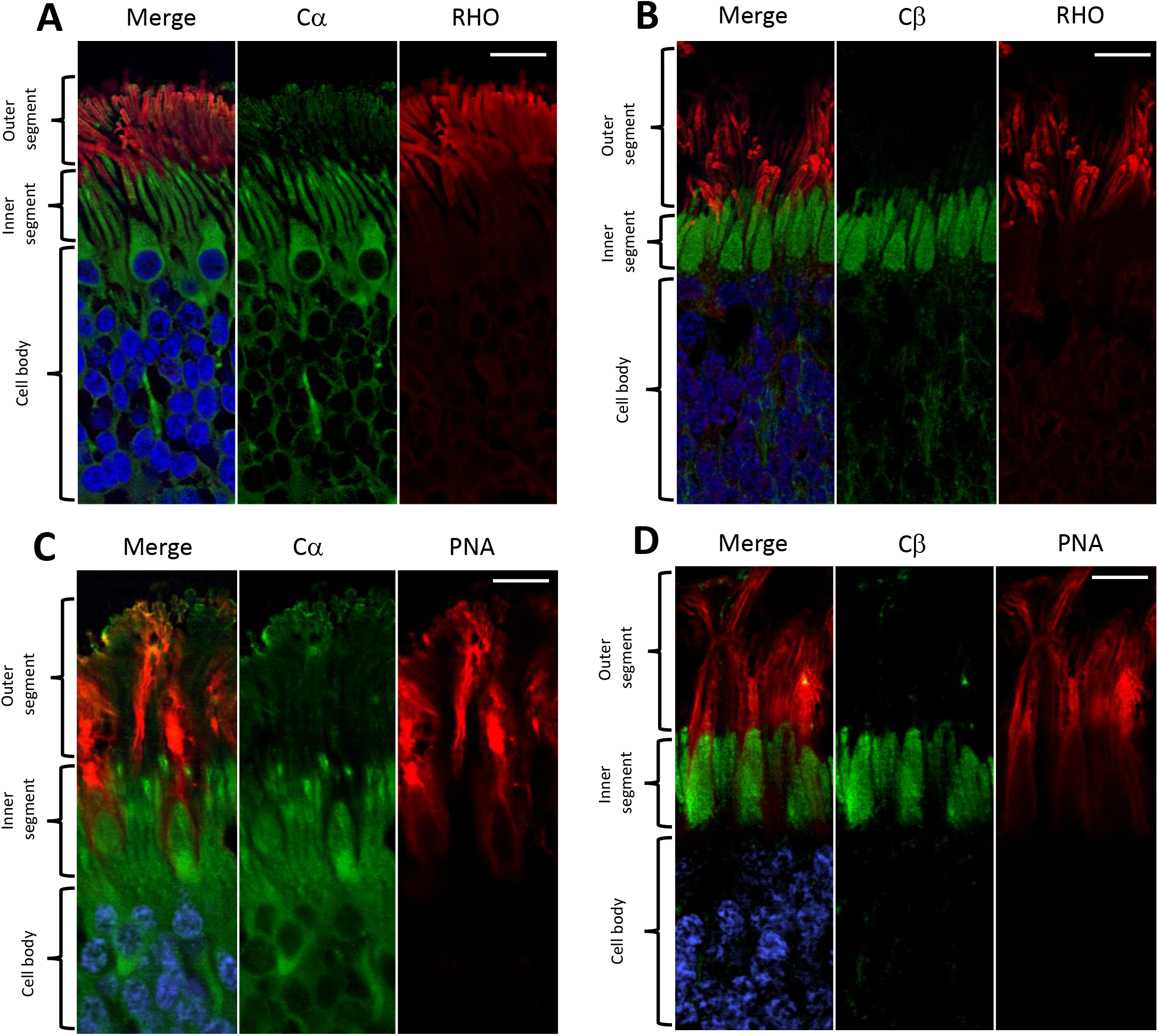
Airyscan images of Cα and Cβ localization in cones and rod cells. Sections stained with anti-Cα or anti-Cβ with anti-RHO (rod cell marker) or anti-PNA (cone cell marker) antibodies highlight intracellular differences in Cα and Cβ localization in rod and cone cells, respectively. **A.** Cα (green) is in the cell body, cilia transition zone, and outer segment membrane, and is distinctly separate from anti-RHO (red) in the outer segment intracellular space. **B.** Cβ (green) is inner segment ellipsoid and is distinctly separate from anti-RHO (red) in the outer segment intracellular space. **C.** Cα (green) is in the cell body, cilia transition zone, and outer segment membrane, and is distinctly separate from anti-PNA (red) in the outer segment intracellular space. **D.** Cβ (green) is inner segment ellipsoid and is distinctly separate from anti-PNA (red) in the outer segment intracellular space.

**Supplemental Figure 2.**
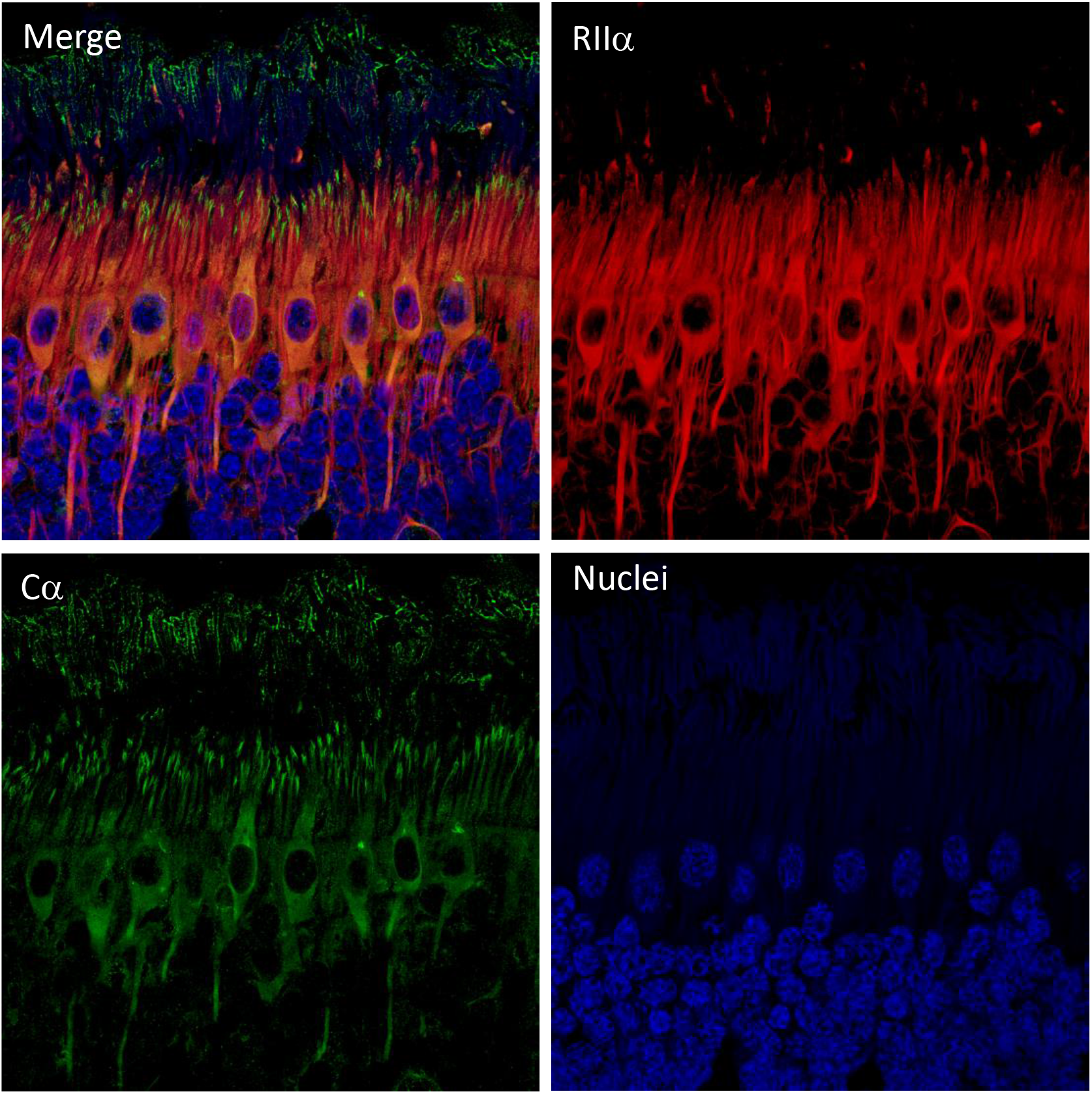
Airyscan images of Cα + RIIα in photoreceptor cells. Z-stack Airyscan images confirm intracellular localization of Cα (green, Cα) and RIIα (red, RIIα), with clear co-localization of Cα and RIIα (yellow, Merge) in the photoreceptor cell body and axon. Only Cα is expressed in the cilia transition zone and outer segment membrane. DAPI stained nuclei are shown in blue.

**Supplemental Figure 3.**
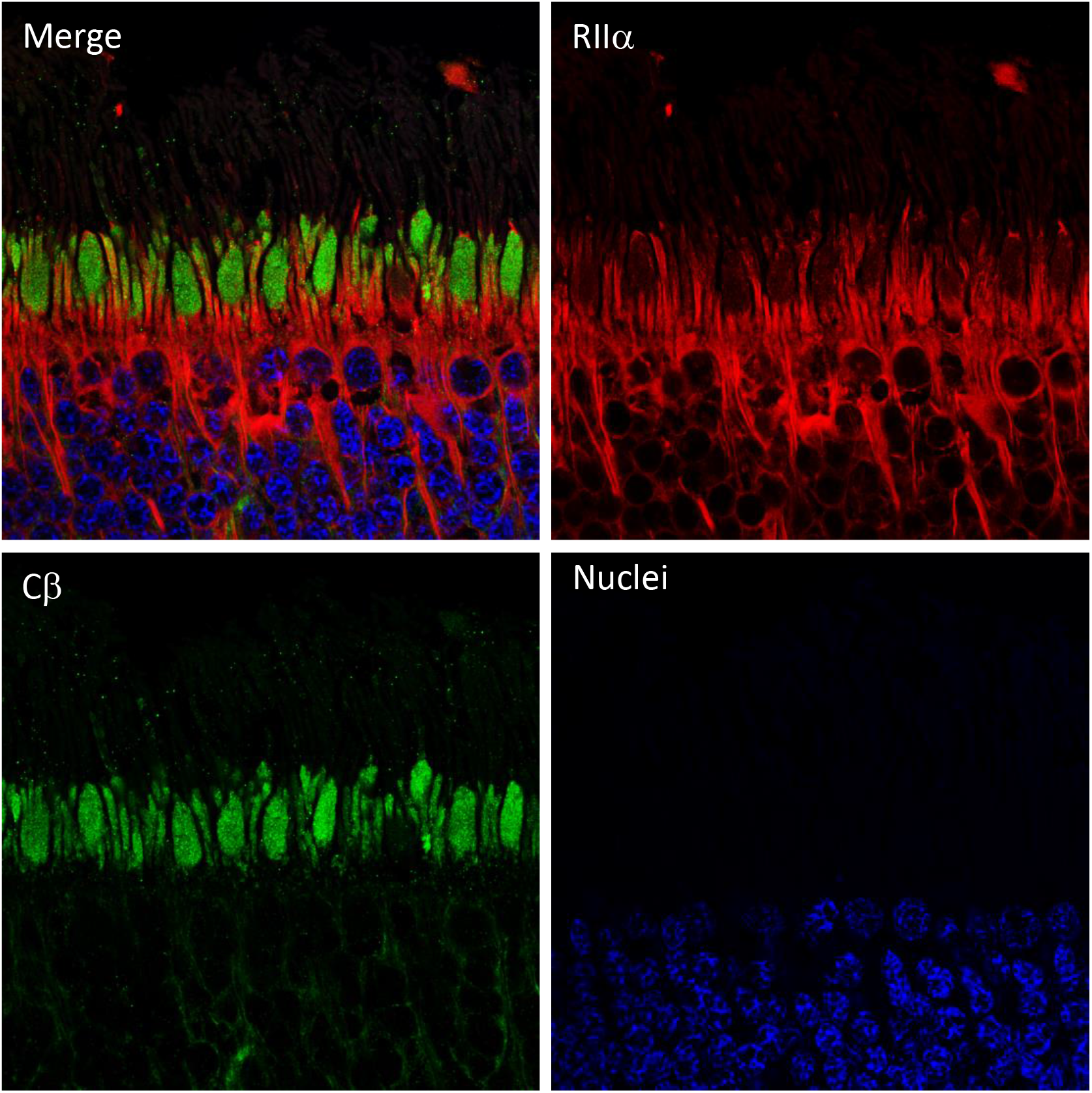
Airyscan images of Cβ + RIIα in photoreceptor cells. Z-stack Airyscan images confirm intracellular localization of Cβ (green, Cβ) and RIIα (red, RIIα), with distinctly different localization of Cβ and RIIα (Merge). Only Cβ is expressed in inner segment ellipsoid, while RIIα is localized to photoreceptor cell body and axon. DAPI stained nuclei are shown in blue.

**Supplemental Figure 4.**
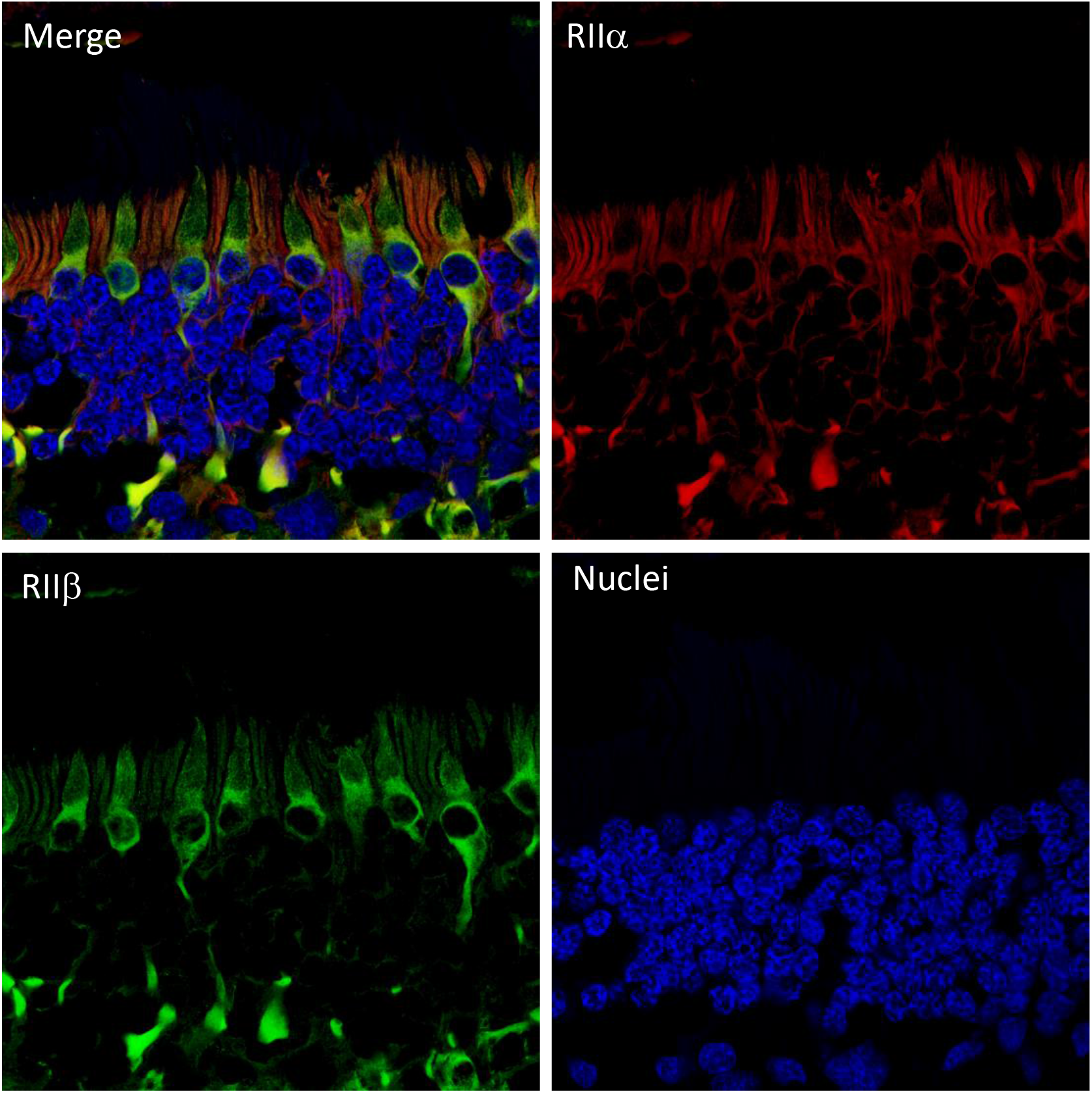
Airyscan images of RIIβ + RIIα in photoreceptor cells. Z-stack Airyscan images confirm intracellular localization of RIIβ (green, RIIβ) and RIIα (red, RIIα), with clear co-localization of RIIβ and RIIα (yellow, Merge) in the cell body and axon of cone cells.

**Supplemental Figure 5.**
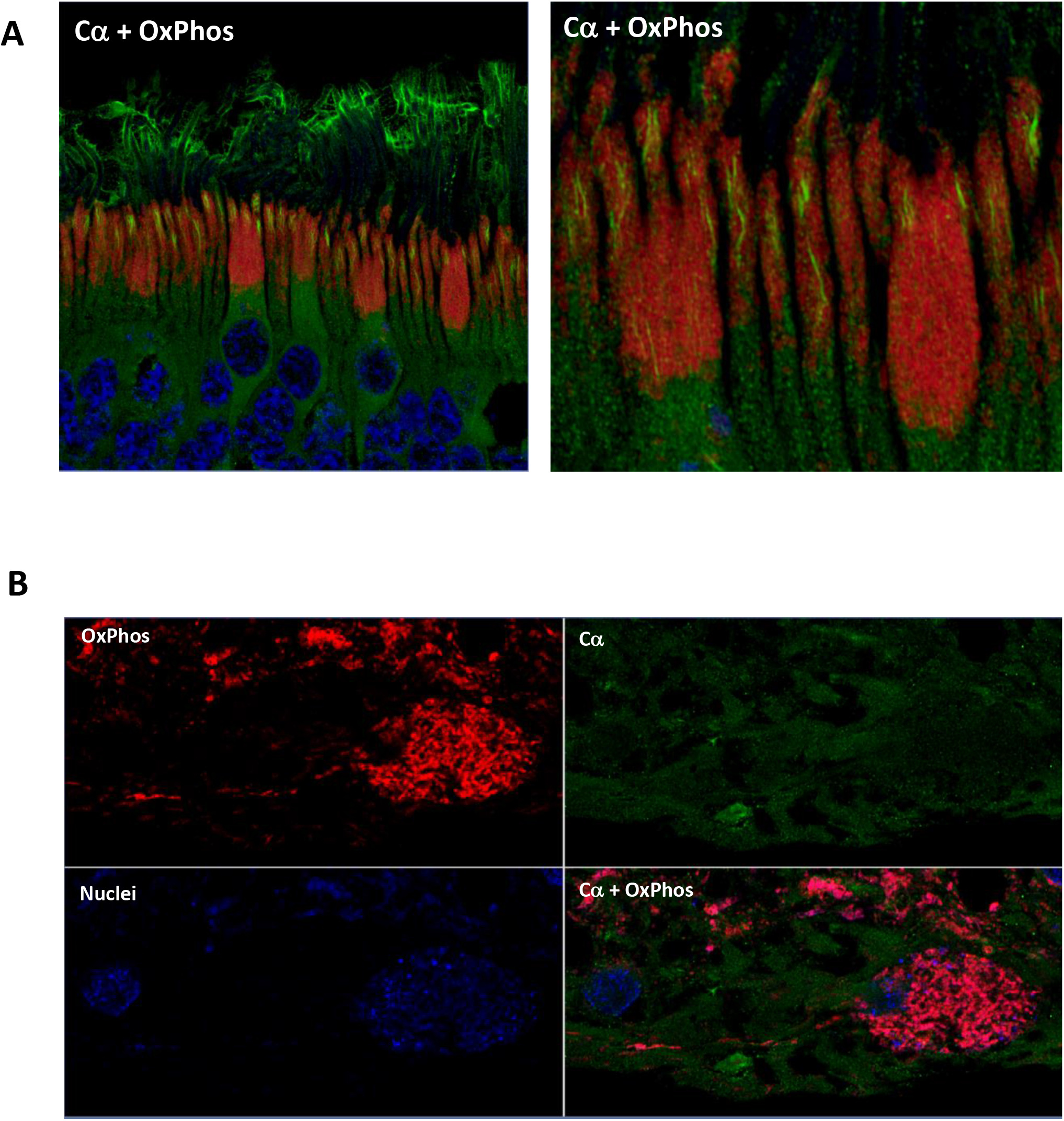
Airyscan images of Cα + mitochondria. Additional z-stack Airyscan images confirm intracellular localization of Cα with respect to mitochondria. **A-B.** Cα (green, Cα) is not localized to mitochondria (red, OxPhos) in photoreceptors (**A**) or retinal ganglion cells (**B**). Nuclei in blue.

**Supplemental Figure 6.**
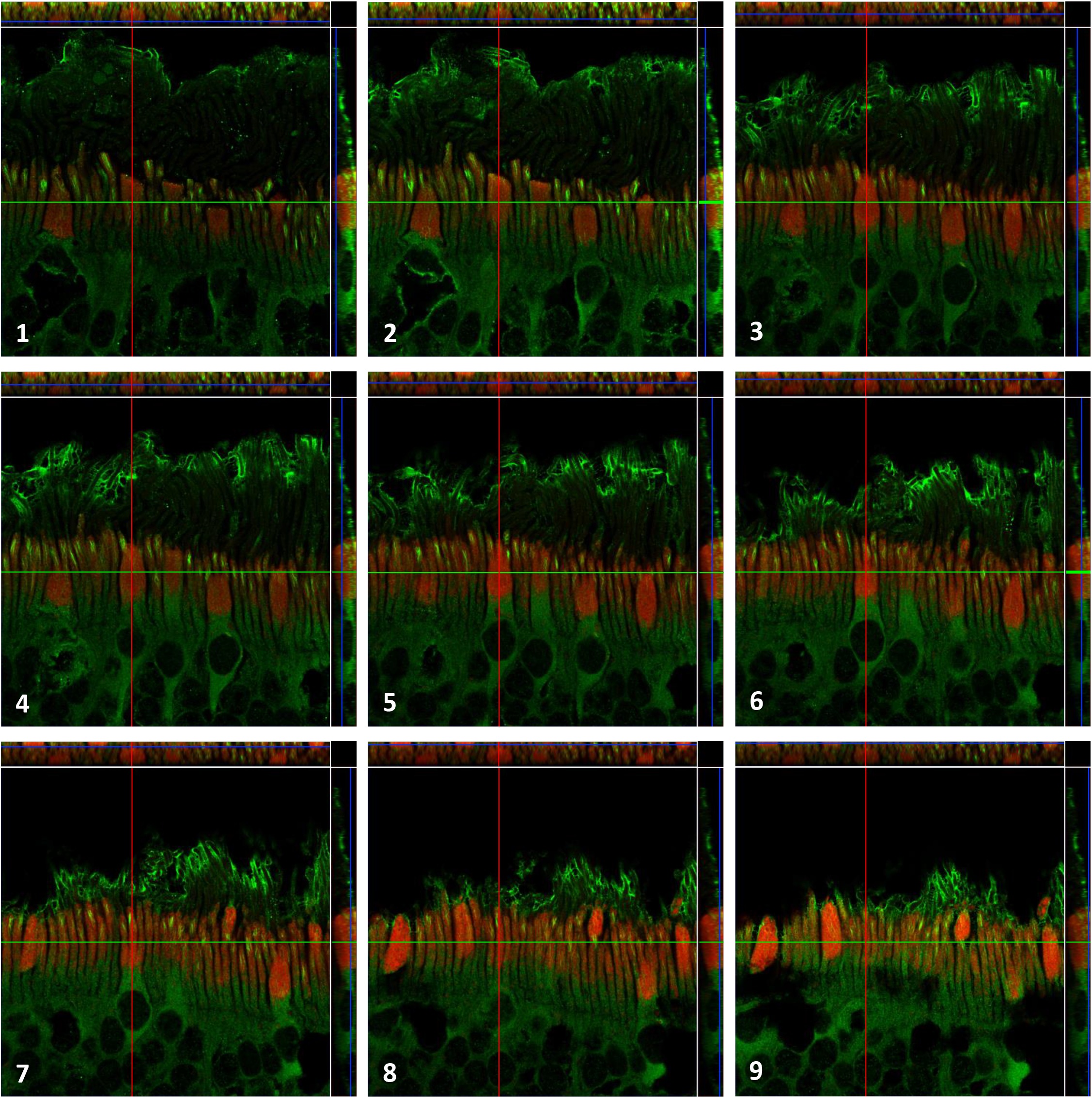
Z-stack images of Cα + mitochondria. Nine z-stack Airyscan images highlight intracellular localization of Cα (green) with respect to mitochondria (red). As the panels move from image 1 (bottom stack, blue line) to image 9 (top stack, blue line) it is clear that Cα is continuously expressed in the outer segment membrane. There is clear signal from Cα (green) at the top of the outer segment in images 1-3, in the mid region in images 4-6, and near the base (closer to the inner segment mitochondria ellipsoid) in images 7-9. Images 1-9 also confirm Cα (green) is not present in the mitochondria ellipsoid (red). Blue line = z position (primary image), green line = top panel, red line = right panel. Green and red lines highlight localization in an individual cell, which can be further visualized in the top and right panel.

**Supplemental Figure 7.**
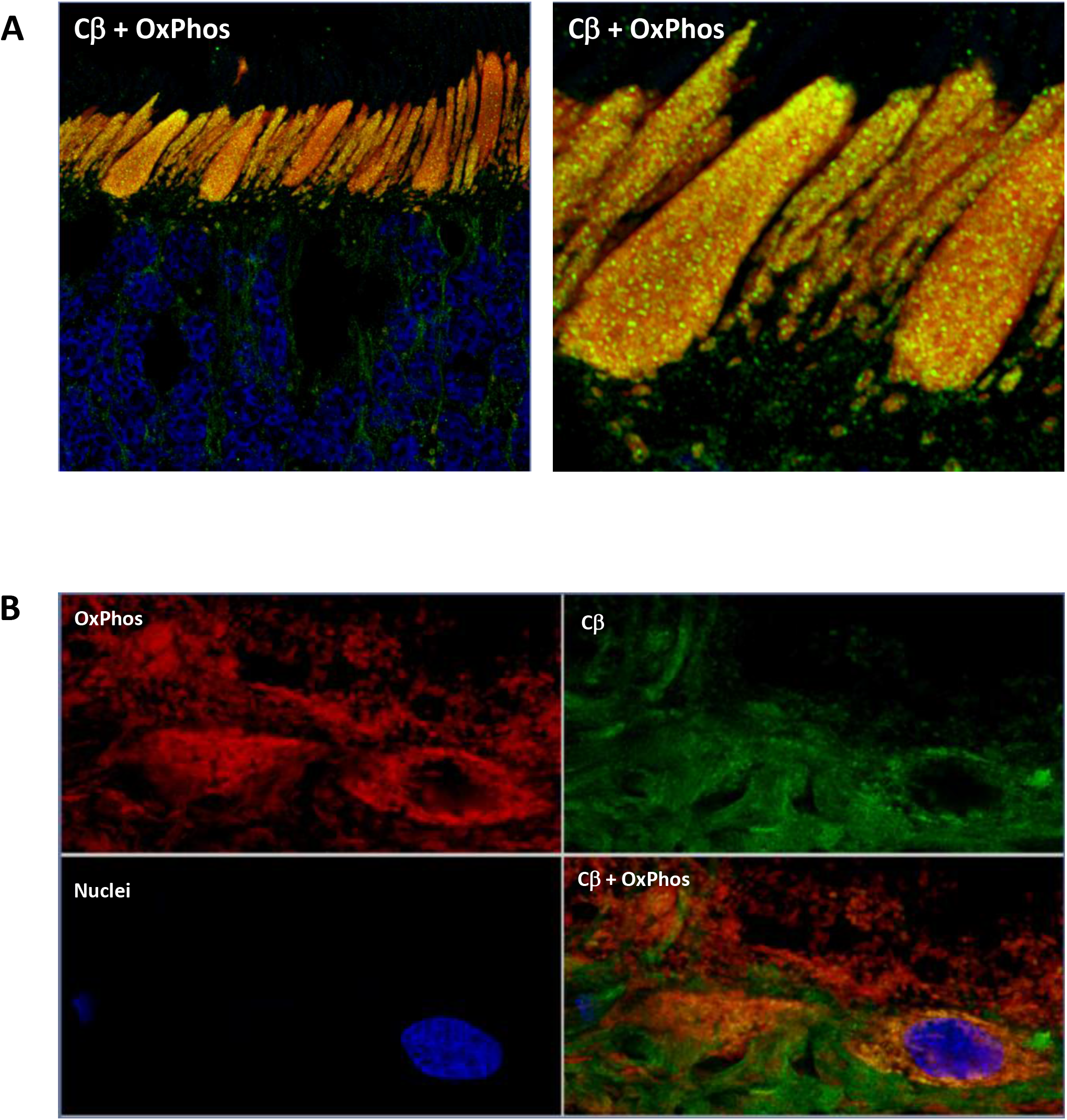
Airyscan images of Cβ + mitochondria. Additional z-stack Airyscan images confirm intracellular localization of Cβ with respect to mitochondria. **A-B.** Cβ (green, Cβ) and mitochondria (red, OxPhos) are clearly co-localized (yellow) in photoreceptors (**A**) and retinal ganglion cells (**B**). Nuclei in blue.

**Supplemental Figure 8.**
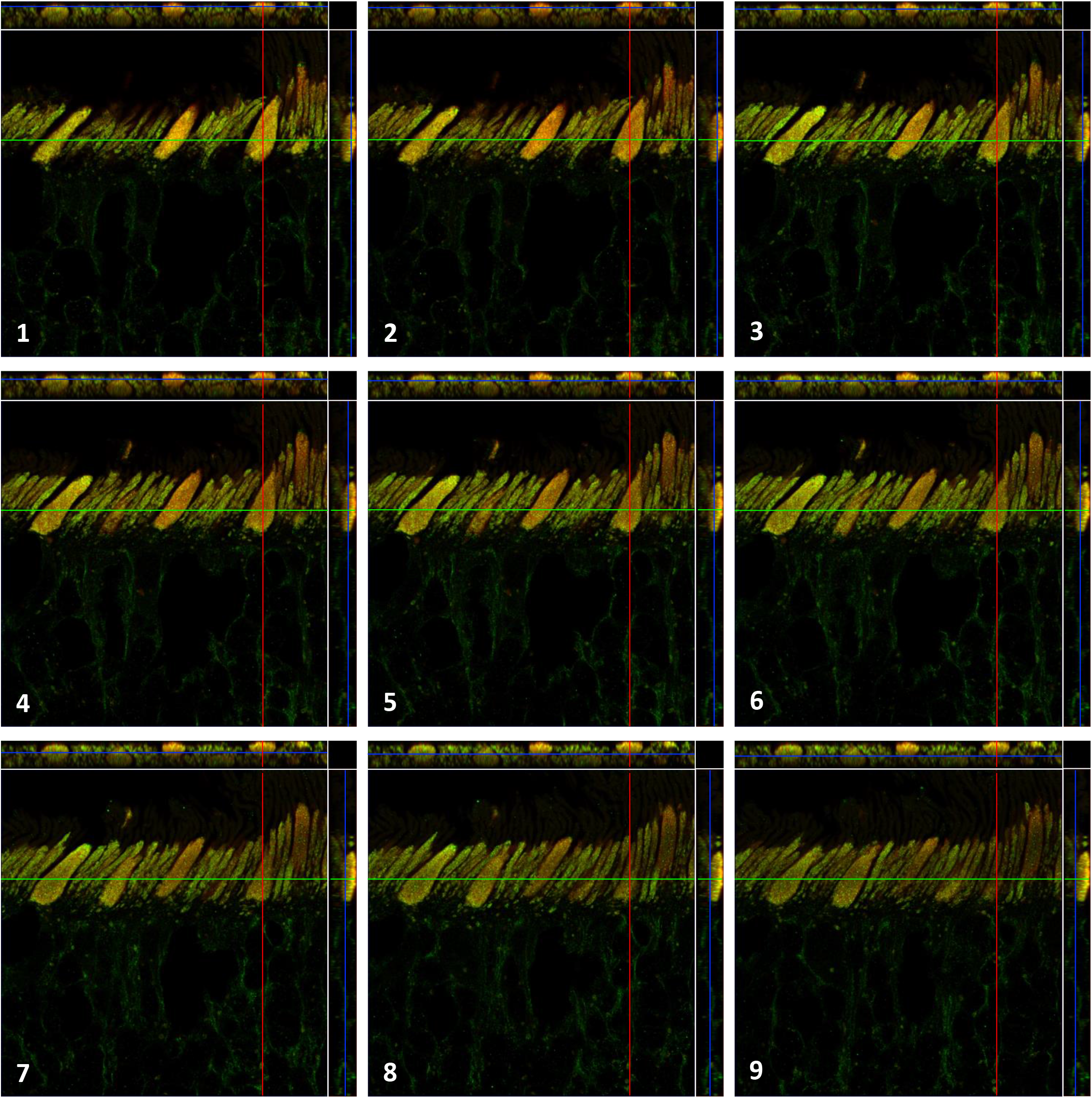
Z-stack images of Cβ + mitochondria. Nine z-stack Airyscan images highlight intracellular localization of Cβ and mitochondria co-localization (yellow). As the panels move from image 1 (top stack, blue line) to image 9 (bottom stack, blue line) it is clear that Cβ is continuously co-localized mitochondria in throughout the entire photoreceptor inner segment. Blue line = z position (primary image), green line = top panel, red line = right panel. Green and red lines highlight localization in an individual cell, which can be further visualized in the top and right panel.

**Supplemental Figure 9.**
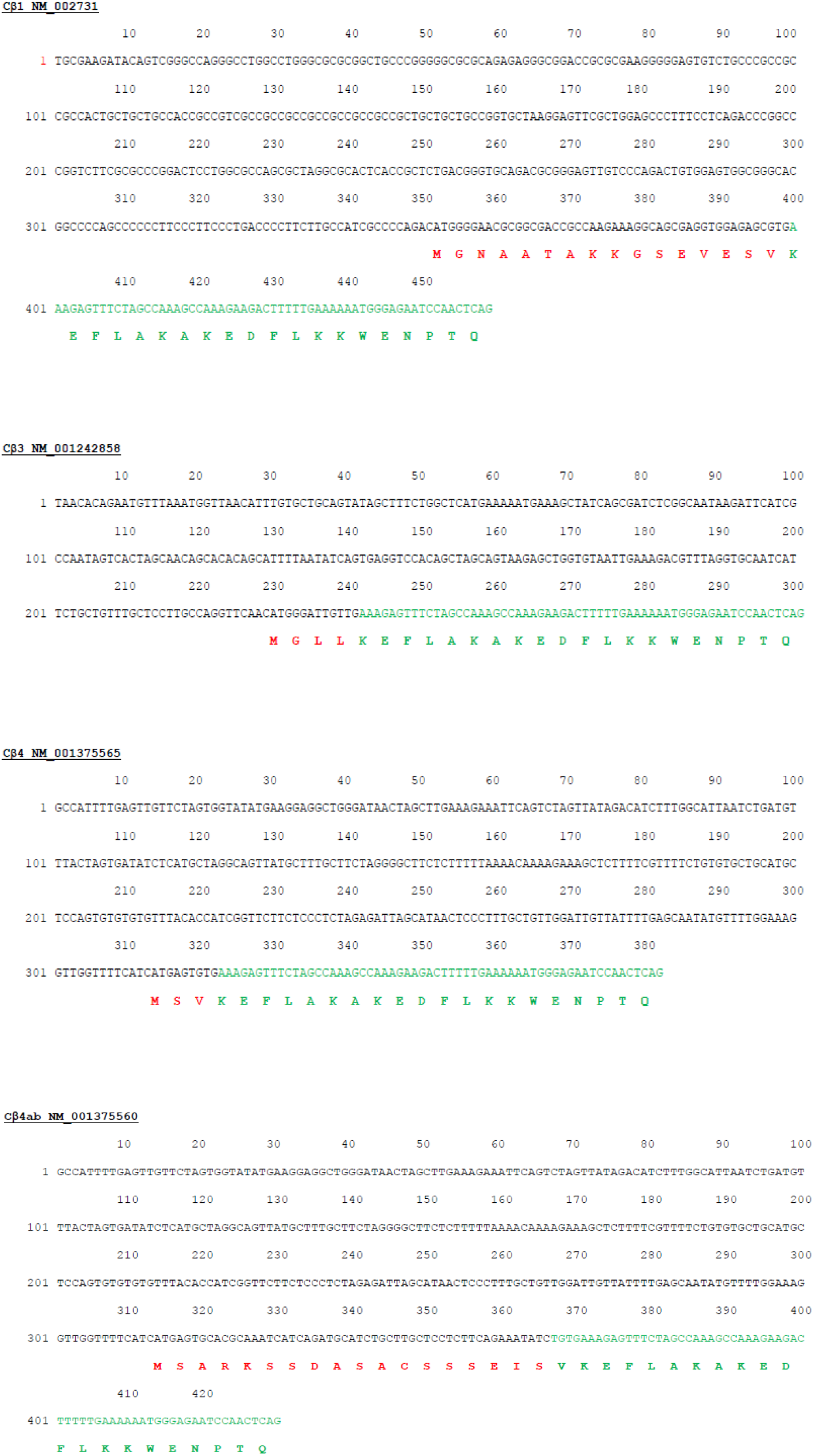
RNA sequence alignments for Cβ1, Cβ3, Cβ4, and Cβ4ab. Differences at the N-terminus were sufficient to design BaseScope probes specific for each isoform (red section), with sequence similarities continuing beyond exon 1 (green section).

**Supplemental Figure 10.**
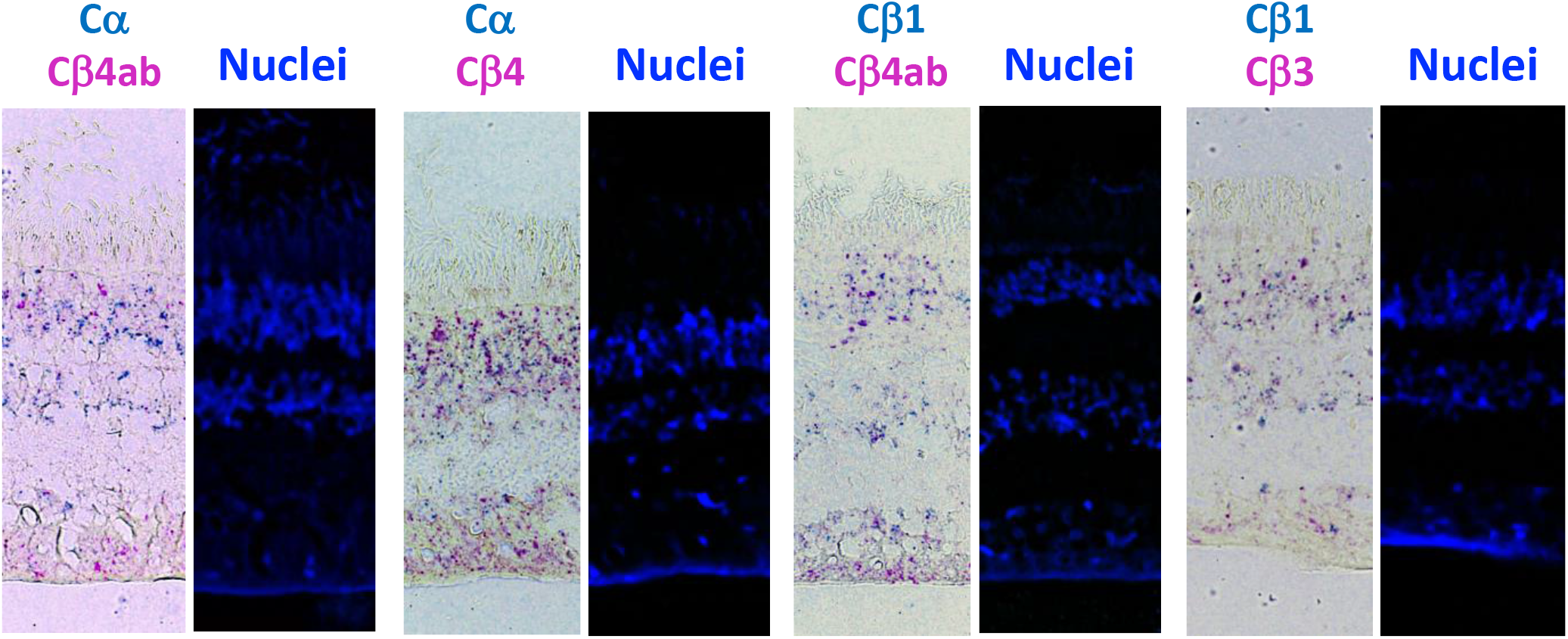
BaseScope Duplex assay of Cα, Cβ1, Cβ3, Cβ4, and Cβ4ab expression. All isoforms are expressed in photoreceptor and interneuron cells, with Cβ4 and Cβ4ab prominently expressed in all tissue layers. Color deconvolution was done following the technical note of ACD (TS 46-003/RevA/Date 6212018). The green color and red color were separated with Image J, darkened, and merged together using Adobe Photoshop.

**Supplemental Figure 11.**
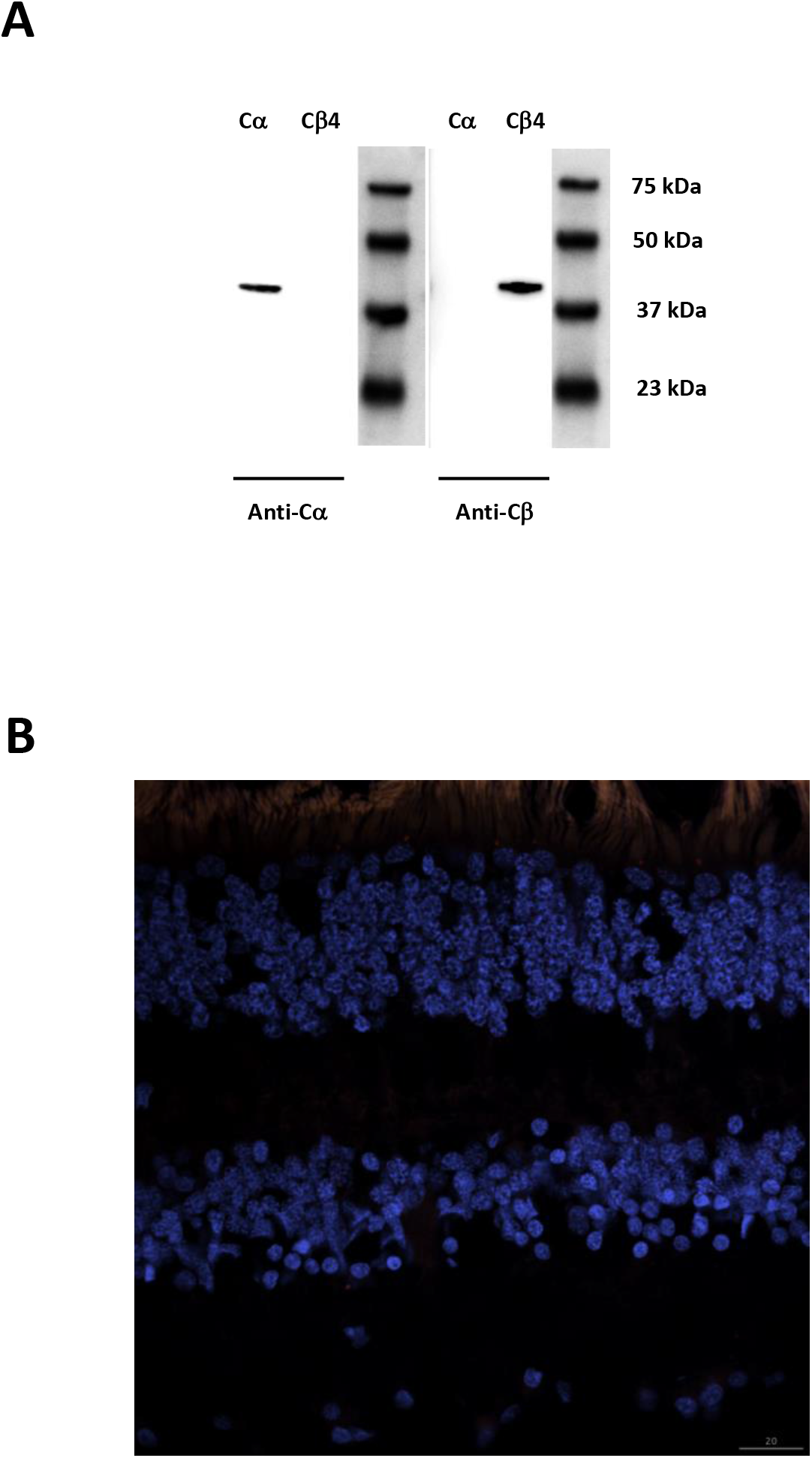
Antibody control for Cα and Cβ. **A.** Anti-Cα antibodies specifically recognized purified Cα protein and not purified Cβ4 protein in Western blots, while anti-Cβ antibodies recognized purified Cα protein and not purified Cβ4 protein. Both Cα and Cβ4 were present at the predicted size of ~39 kDa. **B.** Human retina sections used as negative controls for corresponding IHC data showed no detectable signal after incubation with secondary donkey anti-rabbit and donkey anti-mouse antibodies.

## Notes

### Competing Interest Statement

The authors have declared no competing interest.

